# Extensive *de novo* activity stabilizes epigenetic inheritance of CG methylation in Arabidopsis transposons

**DOI:** 10.1101/2022.04.19.488736

**Authors:** David B. Lyons, Amy Briffa, Shengbo He, Jaemyung Choi, Elizabeth Hollwey, Jack Colicchio, Ian Anderson, Xiaoqi Feng, Martin Howard, Daniel Zilberman

**Affiliations:** Department of Cell & Developmental Biology, John Innes Centre, Norwich, NR4 7UH, UK; Institute of Science and Technology, 3400 Klosterneuburg, Austria; Department of Plant & Microbial Biology, University of California, Berkeley, CA 94720, USA

## Abstract

Cytosine methylation within CG dinucleotides (mCG) can be epigenetically inherited over many generations. Such inheritance is thought to be mediated by a semiconservative mechanism that produces binary present/absent methylation patterns. However, we show here that in *Arabidopsis thaliana h1ddm1* mutants, intermediate heterochromatic mCG is stably inherited across many generations and is quantitatively associated with transposon expression. We develop a mathematical model that estimates the rates of semiconservative maintenance failure and *de novo* methylation at each transposon, demonstrating that mCG can be stably inherited at any level via a dynamic balance of these activities. We find that DRM2 – the core methyltransferase of the RNA-directed DNA methylation pathway – catalyzes most of the heterochromatic *de novo* mCG, with *de novo* rates orders of magnitude higher than previously thought, whereas chromomethylases make smaller contributions. Our results demonstrate that stable epigenetic inheritance of mCG in plant heterochromatin is enabled by extensive *de novo* methylation.

## Introduction

Cytosine methylation provides a mechanism to heritably alter the genome without permanent modification of the DNA sequence (M. Kim & Costello, 2017). DNA methylation represses transposable element (TE) transcription and transposition in plants and vertebrates (H. Zhang et al., 2018). Methylation also regulates endogenous genes: methylation close to the transcriptional start site generally causes gene silencing (Nuñez et al., 2021), whereas methylation of other genic regions can promote or counteract expression (Williams & Gehring, 2020). Genetic defects in the methylation machinery lead to human disease such as cancer (Baylin & Jones, 2016). Disruption of methylation patterns during clonal propagation of plants causes developmental defects that hamper agriculture, and methylation patterns within genes account for a substantial fraction of phenotypic variation in natural *Arabidopsis thaliana* populations (Lloyd & Lister, 2021; Ong-Abdullah et al., 2015; Shahzad et al., 2021). Faithful propagation of DNA methylation is clearly essential, yet our understanding of the underlying processes is far from complete.

Across eukaryotes, cytosine methylation is most prevalent within CG dinucleotides (Schmitz et al., 2019). The core model for the epigenetic inheritance of CG methylation (mCG) – proposed over forty years ago (Holliday & Pugh, 1975; Riggs, 1975) – was inspired by the symmetry of the CG site, and posits that the methylation status of the old strand is used to reproduce the pattern on the strand synthesized during DNA replication. This elegant, semiconservative model has strong experimental support (Catania et al., 2020; Gowher & Jeltsch, 2018; Jeltsch, 2006), but contains a potential flaw because it lacks a mechanism to recover DNA methylation following maintenance failure (Haerter et al., 2014).

Two classes of models have been proposed to address the above issue. The first, developed most explicitly with data from the fungus *Cryptococcus neoformans*, proposes that very high fidelity semiconservative maintenance (estimated failure rate of 9.3×10^−5^ per CG site per cell cycle) combined with rare, random and potentially non-enzymatic *de novo* methylation and natural selection can produce stable epigenetic inheritance of mCG over million-year timescales (Catania et al., 2020). The second, developed with mammalian data, deemphasizes the semiconservative mechanism, and instead proposes a combination of inefficient maintenance (*e.g.* failure rate of 8.0% per site per cell cycle; Wang et al., 2020) and high rates of untemplated *de novo* methylation (*e.g. de novo* rate of 4.7% per site per cell cycle; Wang et al., 2020) that is specifically targeted to methylated regions by an unknown mechanism (Haggerty et al., 2021; Lövkvist et al., 2016; Wang et al., 2020). The maintenance methyltransferases in the two systems are different (Dnmt5 in *C. neoformans* and Dnmt1 in mammals; Dumesic et al., 2020; Huff & Zilberman, 2014), which might account for the distinct mechanisms. The overall lack of long-term, transgenerational epigenetic inheritance of mammalian mCG (Kazachenka et al., 2018; Seisenberger et al., 2013; Tucci et al., 2019) may also be compatible with higher rates of maintenance failure.

The existing plant data (mostly from *Arabidopsis*) appear to be more compatible with the former mechanism. Plants exhibit stable, transgenerational mCG inheritance (Becker et al., 2011; Quadrana & Colot, 2016; Schmitz et al., 2011), and although plants use MET1 (a Dnmt1 family enzyme; Tirot et al., 2021), the global maintenance failure rate of 6.3×10^−4^ per generation (Van Der Graaf et al., 2015) or about 1.9×10^−5^ per cell cycle (assuming 34 cell cycles per generation, Watson et al., 2016) reported for *Arabidopsis* is lower than that reported for *C. neoformans* (9.3×10^−5^, Catania et al., 2020). Even the higher maintenance failure rate of 4.4×10^−5^ per cell cycle (1.5×10^−3^ per generation) reported for *Arabidopsis* genes (Van Der Graaf et al., 2015) is two times lower than the *C. neoformans* rate. Epigenetic inheritance of plant mCG is thought to depend only on MET1 (Cokus et al., 2008; Lister et al., 2008; H. Zhang et al., 2018), which, like *C. neoformans* Dnmt5, does not have known *de novo* activity (H. Zhang et al., 2018). The reported rates of *de novo* mCG are also low in *Arabidopsis* (global rate of 2.6×10^−4^ per generation or 7.6×10^−6^ per cell cycle; Van Der Graaf et al., 2015), and the source of the *de novo* activity is unknown.

Plant and animal genomes also contain cytosine methylation outside CG dinucleotides (Du et al., 2015). In plants, the CMT3 methyltransferase family catalyzes methylation of CNG trinucleotides, conventionally described as CHG (where H is any non-G base) to avoid overlapping CG sites (Law & Jacobsen, 2010). The related CMT2 family can methylate cytosines outside CG and CNG contexts (Zemach et al. 2013; Stroud et al. 2014), a pattern of specificity referred to as CHH. CMT2 and CMT3 rely on a positive feedback loop with dimethylation of lysine 9 of histone H3 (H3K9me2) (Du et al., 2012, 2015; Rajakumara et al., 2011) – a hallmark of heterochromatin (Sequeira-Mendes et al., 2014) – and therefore preferentially methylate heterochromatic TEs (Stroud et al., 2014; Zemach et al., 2013; Y. Zhang et al., 2018). Plants also possess an RNA-directed DNA methylation (RdDM) pathway, in which 24 nucleotide RNA molecules guide DRM methyltransferases (homologs of animal Dnmt3) to initiate DNA methylation in all sequence contexts, and to maintain CHH methylation at relatively euchromatic TEs (Matzke & Mosher, 2014).

Regardless of the mechanism, plants do not perfectly copy non-CG methylation each cell division, which leads to a probabilistic distribution of methylation states at CHG and CHH sites that contrasts with the more binary mCG patterns (Cokus et al., 2008; Lister et al., 2008). Although the RdDM pathway can establish DNA methylation in every context, including CG (Chan et al., 2004; Cuerda-Gil & Slotkin, 2016; Pélissier et al., 1999; Teixeira et al., 2009), several studies have concluded that mCG maintenance is independent of the non-CG methylation pathways (Lister et al., 2008; Stroud et al., 2014; To et al., 2022). The *de novo* mCG rates reported for TEs (where the non-CG pathways operate) are also lower than those for genes (Van Der Graaf et al., 2015), ranging down to 6.8×10^−8^ per cell cycle (2.3×10^−6^ per generation; Hazarika et al., 2022). Thus, inheritance of mCG within TEs is thought to be essentially semiconservative.

Previously, we described widespread intermediate mCG within heterochromatic TEs of *Arabidopsis h1dddm1* mutants lacking the nucleosome remodeler DDM1 and linker histone H1 (Zemach et al. 2013; Lyons et al. 2017). The reduced mCG maintenance efficiency in *h1ddm1* mutants combined with the reported low *de novo* rates in TEs (Hazarika et al., 2022; Van Der Graaf et al., 2015) imply that these intermediate mCG patterns should be unstable and eventually degrade to very low levels, but this has not been evaluated. Here we analyze *h1ddm1* DNA methylation over ten generations of inbreeding. We find that, contrary to the above expectation, intermediate mCG patterns are stably inherited. Using a mathematical model, we find a predicted *de novo* component of mCG inheritance orders of magnitude stronger than implied by the previously reported transgenerational rates of mCG gain in TEs. This prediction is confirmed by analysis of *h1ddm1* compound mutants lacking either CMT2, CMT3, or the key RdDM methyltransferase DRM2. Our results contradict the established view that mCG epigenetic inheritance within TEs is essentially semiconservative. Instead, our data indicate that epigenetic inheritance of heterochromatic mCG is stabilized by extensive *de novo* methylation, with RdDM contributing most of the *de novo* activity and CMT2/3 making smaller contributions.

## Results

### TE CG methylation decays to stable intermediate levels within ten *h1ddm1* generations

To understand DNA methylation inheritance in *h1ddm1* plants, we crossed a plant homozygous for mutations in both canonical *Arabidopsis* H1 genes to a heterozygous *ddm1* plant from a line in which the *ddm1* mutation had never been homozygous. The resulting *h1.1/+;h1.2/+;ddm1/+* F1 was allowed to self-fertilize to generate an F2 founder in which all three mutant alleles were homozygous (and the *ddm1* allele was homozygous for the first time). As DDM1 predominantly functions in heterochromatin (Zemach et al., 2013), we focused our analysis on heterochromatic TEs (hTEs) that lose much of their DNA methylation in all sequence contexts when this remodeler is inactivated (Fig. 1A-C and Fig. S1A-B). Consistent with the positive association between DNA methylation loss and H3K9me2 in *ddm1* plants (Zemach et al., 2013), F2 *h1ddm1* mCG varies substantially between hTEs and shows a strong negative correlation with H3K9me2 in the parental *h1* genotype (r = −0.60, Fig. 1D). Unlike *ddm1* where most mCG is lost by the F2 (Fig. S1C), average and median *h1ddm1* mCG across hTEs decreases substantially from the F2 to the F3 generation (median change 11%), less so from the F3 to the F4 generation (~5%), and even less from the F4 to the F5 (Fig. 1A-C). Overall mCG remains stable at intermediate levels after the F5 generation, up to generation F11 (ten generations of inbreeding; Fig. 1A-C, Fig. S1D). Across all ten *h1ddm1* generations analyzed, intermediate mCG spans vast swaths of heterochromatin (>12Mbp, ~10% of the genome), as can be appreciated from genome browser views (Fig. 1B). CG methylation patterns are well-correlated across generations (Fig. 1E), indicating that the heterogeneous mCG landscape of *h1dddm1* hTEs is accurately reproduced.

**Figure 1.**
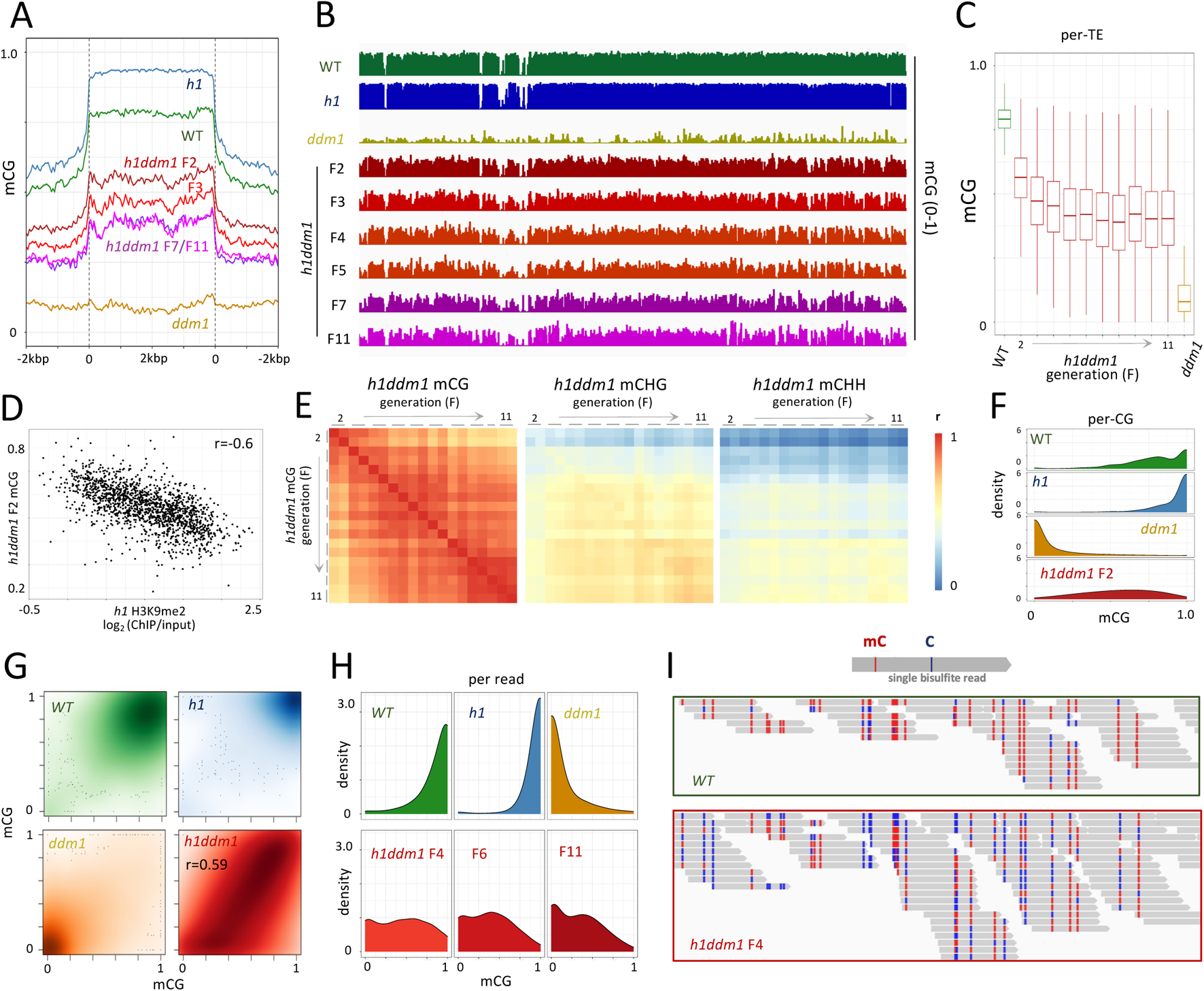
Heterochromatic mCG decays to an intermediate equilibrium in *h1ddm1*. **A)** Average fractional CG methylation (mCG) is plotted for all heterochromatic TEs >250 bp (Choi et al., 2020) in the indicated genotypes and *h1ddm1* generations. **B)** Genome browser view of a 400 kb window of pericentromeric heterochromatin illustrating mCG across *h1ddm1* generations (Chr. 5:12,950,000-13,350,000). **C)** Boxplots of mCG for all *h1ddm1* heterochromatic TEs (red), beginning with *WT* (green, left) and ending with *ddm1* F5 (dark orange, right). Box depicts the interquartile range and whiskers extend to 1.5X interquartile range; horizontal line shows the median value. **D)** F2 *h1ddm1* per-TE mCG for all TEs >2 kb as a function of H3K9me2 (expressed as log_2_ ChIP/input) in the parental genotype, *h1*. **E)** Heatmaps of per-TE mCG Pearson’s correlation (color bar) for mCG vs. mCG, mCHG, and mCHH across *h1ddm1* generations (x-axes, from left to right respectively). Biological replicates are indicated with a grey bar over columns/rows (one replicate for the F10, two for all other generations). Note increased correlation between CG and non-CG methylation in later generations. **F)** Density plot of per-CG methylation for all CG sites with coverage >10 in heterochromatic TEs for the indicated genotypes. **G)** 2D density plots of per-CG methylation for all CG sites in heterochromatic TEs for F4 vs. F5 generations (except for *h1ddm1*, where F2 vs. F3 is shown to illustrate the correlation between intermediate values across generations at a stage when mCG is changing the most). Darker color indicates more data points. **H)** Density plots of mCG per-bisulfite-read with >=3 CG sites per read from heterochromatic TEs. **I)** Genome browser view of bisulfite reads for *WT* (top) and *h1ddm1* (bottom); Chr.1:14,918,800-14,918,900. Only (+) strand mapping reads are shown for clarity.

CHG methylation (mCHG) behaves similarly to mCG over *h1ddm1* generations, dropping substantially initially, then leveling off in the subsequent generations (Fig. S1A,E), whereas *h1ddm1* CHH methylation (mCHH) decreases overall, though not monotonically (Fig. S1B,F). In both cases relative losses are much smaller than for mCG. Early generation mCHG and mCHH patterns are poorly correlated with mCG, but the correlation improves over time (Fig. 1E), suggesting that methylation in all contexts evolves in concert across *h1ddm1* generations. Although correlations are strongest between methylation patterns of the late generations, early generation mCHG and mCHH patterns are better correlated with late generation mCG than with early generation mCG (Fig. 1E). Therefore, to some extent, early generation non-CG methylation can predict late generation mCG. This observation suggests that the processes shaping mCG epigenetic inheritance in TEs are related to non-CG methylation.

### Intermediate mCG is caused by methylation fluctuations at individual CG sites

A simple explanation for the observed intermediate mCG across *h1ddm1* generations would be methylation heterogeneity across CG sites, with fully methylated and fully unmethylated sites producing an intermediate pattern when averaged in bins. In this scenario, the gradual decrease and stabilization of mCG in *h1ddm1* would be caused by an increasing number of CG sites permanently switching to an unmethylated state, until only sites with stable mCG maintenance remain methylated around the F5 generation. For example, because DDM1 is more important for mCG maintenance in nucleosomes than in the connecting linker DNA (Lyons & Zilberman, 2017), the observed intermediate mCG patterns could be caused by stable mCG maintenance in linkers and unstable maintenance in nucleosomes. This should cause mCG in TEs with poorly positioned nucleosomes to stabilize at lower levels, because shifting nucleosome positions across cell cycles should prevent efficient maintenance at any CG site. However, this is not the case, as mCG behaves similarly across *h1ddm1* generations in TEs with various levels of nucleosome positioning (Fig. S1G).

To test the broader possibility of bimodal CG site methylation heterogeneity, we plotted per-base methylation in wild type (*WT*), *h1, ddm1* and *h1ddm1* plants for all heterochromatic CG sites methylated above 5% in *WT*. The near-binary nature of mCG is evident in *WT* versus *ddm1* plants, and even more so in *h1* versus *ddm1* mutants (Fig. 1F), because *h1* enhances heterochromatic mCG (Fig. 1A-B; Lyons & Zilberman, 2017; Zemach et al., 2013). In contrast, F2 *h1ddm1* hTE mCG is much more uniformly distributed, covering with substantial representation all methylation values from 0 to 1 (Fig. 1F). This pattern largely persists over the generations (Fig. S1H) and is correlated across generations (Fig. 1G and S1I). Therefore, CG sites that are typically consistently methylated across a *WT* population of cells exist in a mixture of methylation states in *h1ddm1* cells.

Another possibility to generate intermediate mCG is methylation heterogeneity across entire loci (TEs, for example), with fully methylated and fully unmethylated loci producing an intermediate pattern when averaged. This scenario entails dynamic switching of locus methylation, with the fraction of loci that lose mCG increasing over the first few generations before stabilizing. To examine this possibility, we analyzed methylation of sequencing reads that correspond to individual DNA molecules, where locus-level variation should produce reads with either low or high methylation. When considering all heterochromatic reads with >3 CG dinucleotides in early, middle, and late generations, the observed distribution of per-read mCG is relatively uniform in *h1ddm1* plants (Fig. 1H), similar to the per-CG *h1ddm1* mCG (Fig. 1F), indicating that many individual DNA molecules have a mixture of methylated and unmethylated CG sites. This becomes apparent when mCG patterns of individual reads are visualized, with *h1ddm1* reads exhibiting a mixture of mCG levels and patterns (Fig. 1I). These results demonstrate that the intermediate *h1ddm1* mCG patterns do not arise from averaging of fully methylated and unmethylated CG sites, or from entire loci dynamically switching their methylation states during development. Instead, our results indicate that individual CG sites regularly switch their methylation states, implying a substantial flux of *de novo* mCG in *h1ddm1*.

Although *WT* CG sites tend to have higher methylation levels than in *h1ddm1*, individual CG sites nonetheless show methylation variability between DNA molecules in *WT* (Fig. 1I). This is reflected in a broader distribution of per-read methylation levels in *WT* compared to *h1* (Fig. 1H) and in a similarly broader distribution of per-CG methylation (Fig. 1F-G). This suggests that mCG epigenetic inheritance in *WT* heterochromatin also involves substantial *de novo* methylation.

### Intermediate CG methylation is heritable in *h1ddm1* plants

A possible explanation for the apparently stable inheritance of intermediate heterochromatic mCG in *h1ddm1* plants is that we are assaying the wrong tissue. We measure DNA methylation in leaves, which do not contribute to the subsequent generation. Cells that mediate inheritance, such as reproductive cells, show more robust maintenance of mCG than leaves and other somatic tissues (Hsieh et al., 2016; Park et al., 2016). Therefore, the *h1ddm1* genotype might primarily destabilize somatic, but not reproductive methylation maintenance. In this scenario, the intermediate mCG patterns we observe in leaves may be produced in each generation through methylation decay during somatic development from an initially high level expected to be compatible with primarily semiconservative inheritance. To test this hypothesis, we quantified sperm DNA methylation in the F4 and F5 generations of *h1ddm1* plants and *WT*, *h1* (only the F5 generation), and *ddm1* siblings. *WT* plants have increased mCG in sperm compared to leaf (Fig. 2A-B), in agreement with previous work (Hsieh et al., 2016), whereas mCG is similarly low in *ddm1* sperm and leaf (Fig. 2A,C). Sperm mCG is unchanged in *h1* compared to *WT* (Fig. 2A-C), consistent with the low H1 levels in male reproductive cells (He et al., 2019). Additionally, we find a large reduction of mCHH (but not mCHG) in sperm compared to leaf in all genotypes (Fig. S2A-B), indicating that mCHH reprogramming in sperm (Calarco et al., 2012) does not require DDM1.

**Figure 2.**
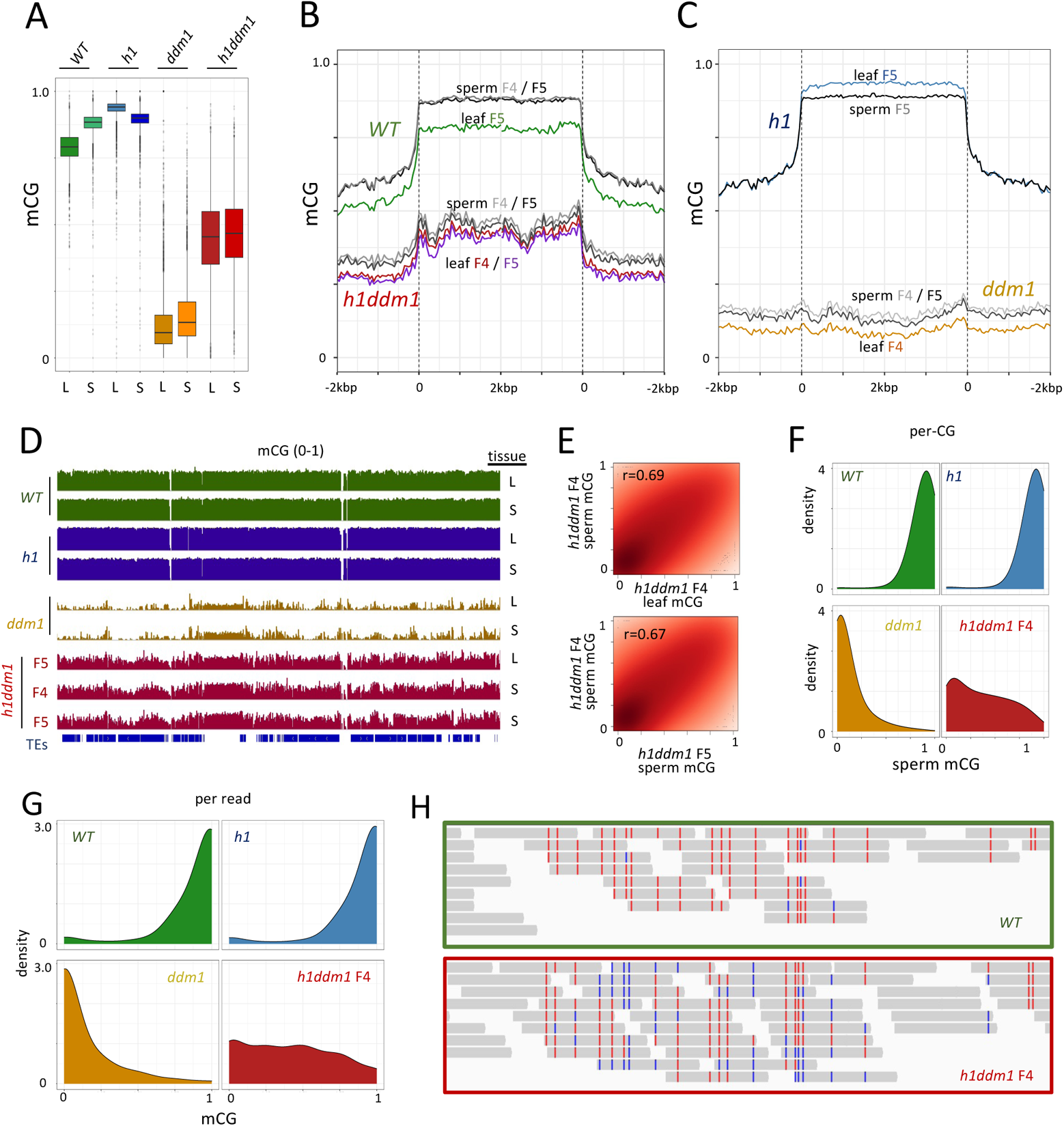
Intermediate *h1ddm1* mCG is heritable. **A)** Boxplots of F5 mCG for all heterochromatic TEs with leaf values (L) displayed next to sperm (S) in the indicated genotypes. **B-C**) Average mCG in sperm and leaf for the indicated generations and genotypes plotted relative to the start and end sites of heterochromatic TEs, as in Fig. 1A. **D)** Genome browser view of 500 kb of heterochromatin (Chr 5: 11,050,000-11,550,000) highlighting the similarities between sperm and leaf *h1ddm1* mCG. Source tissue is either leaf (L) or sperm (S). All non-*h1ddm1* genotypes shown are F5 generation. **E)** 2D density plots comparing per-CG methylation at sites with coverage >10 in F4 sperm vs. F4 leaf (top) or F5 sperm (bottom) across heterochromatic loci. Pearson’s r is shown. Darker color indicates more data points. **F)** Density plots of sperm per-CG methylation for all CG sites with coverage >5 in heterochromatic TEs. **G)** Density plots of sperm mCG per-bisulfite-read with ≥3 CG sites per read. **H)** Screenshot of per-read methylation from F4 sperm of *WT* and *h1ddm1* at Chr.3: 12,425,889-12,426,499. Only (+) strand is shown for clarity.

Importantly, *h1ddm1* plants possess nearly the same average mCG in sperm and leaf in the F4 and the F5 generations (Fig. 2A-B), a phenomenon readily apparent in the genome browser (Fig. 2D). Furthermore, there is a high correlation of per-site mCG between sperm and leaf of the same generation (Fig. 2E, top) as well as sperm across generations (Fig. 2E, bottom). Similar to *h1ddm1* leaves (Fig. 1F), *h1ddm1* sperm per-site mCG shows a continuous distribution of methylation frequencies (Fig. 2F). Per-read sperm methylation patterns in *h1ddm1* (Fig. 2G-H) are also similar to those of leaves (Fig. 1H-I), indicating a mixture of mCG levels and patterns in *h1ddm1* sperm cells.

Because sperm cells directly initiate the next generation, our results demonstrate that patterns of intermediate methylation are indeed inherited. The extensive mCG heterogeneity between DNA molecules (Fig. 2G-H), which represents heterogeneity between the haploid sperm, indicates that widespread *de novo* methylation of CG sites mediates stable inheritance of heterochromatic mCG in *h1ddm1* plants. Notably, per-read mCG heterogeneity is also apparent in *WT* sperm (Fig. 2G-H), suggesting an important role for *de novo* methylation in maintaining *WT* heterochromatic mCG.

### TSS methylation is quantitively associated with TE expression in *ddm1* and *h1ddm1* plants

Because loss of DNA methylation is accompanied by widespread TE activation in *ddm1* mutants (Miura et al., 2001; Tsukahara et al., 2009), we examined whether the increased levels of DNA methylation in *h1ddm1* (compared to *ddm1*) are associated with reduced TE expression. After assembling transcripts based on the Araport11 assembly (Pertea et al., 2015), which includes *ddm1*-specific TE transcripts (Panda & Slotkin, 2020), we quantified expression in *WT*, *h1*, *ddm1* and *h1ddm1* plants. Of the 1375 transcripts upregulated in *ddm1*, 1075 overlap heterochromatic TEs. 796 of these are bona fide expressed heterochromatic loci, based on their H3K9 methylation profiles and expression level (H3K9me1 or -me2 > 0.25 in *WT*, >2.5 TPM in *ddm1*; Fig. S3A). Expression of these heterochromatic transcripts is generally reduced in *h1ddm1* compared to *ddm1* (Fig. 3A-B), with 303 of these loci significantly downregulated (Fig. S3B, qval<0.05). In contrast, expression of upregulated euchromatic *ddm1* transcripts is not substantially altered in *h1ddm1* (Fig. S3C).

**Figure 3.**
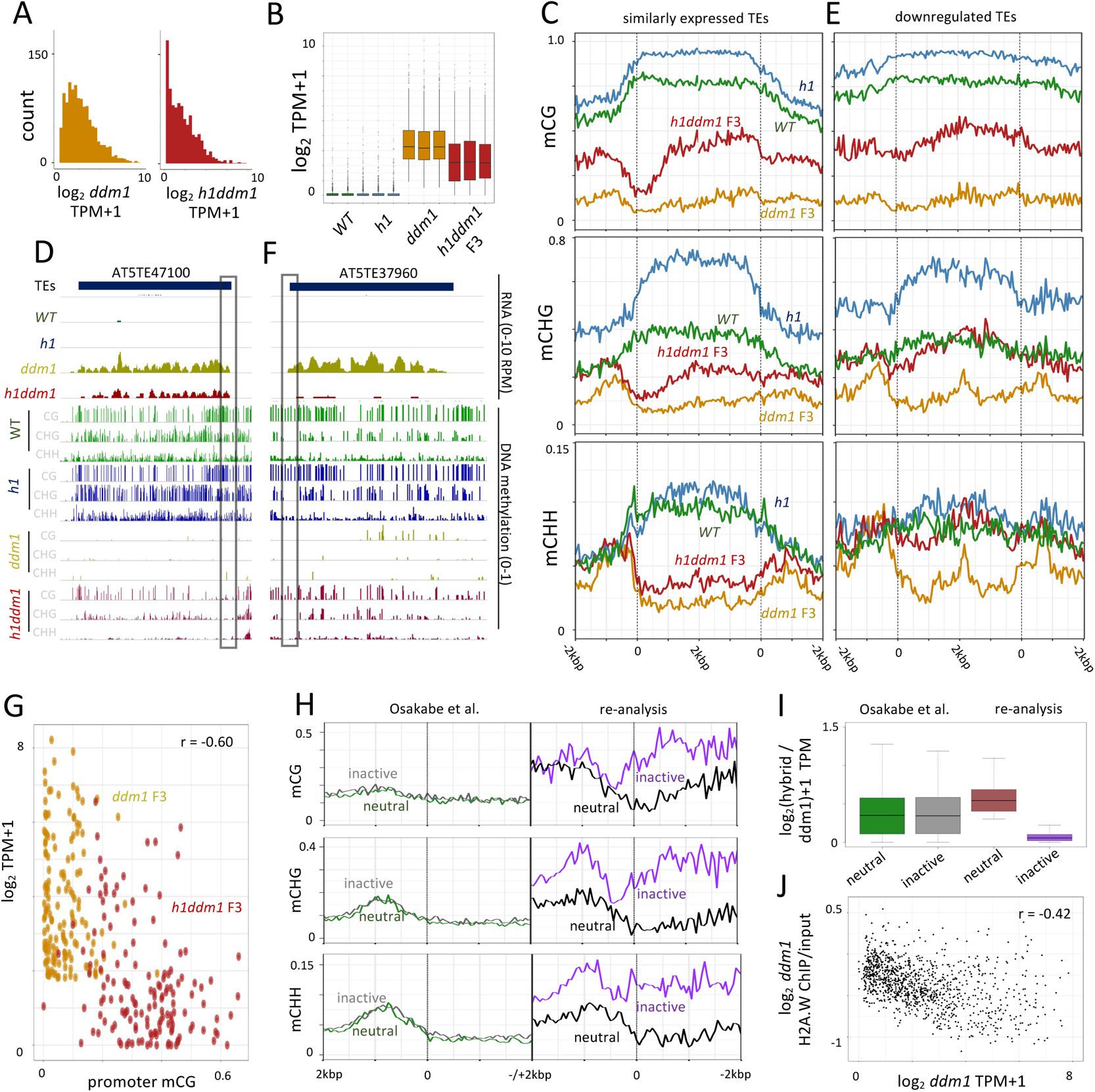
Promoter methylation is quantitatively associated with TE expression. **A)** Histograms of replicate-averaged expression levels (log_2_ transformed transcript-per-million, TPM) of *ddm1* de-repressed heterochromatic transcripts with expression above 2.5 TPM (n=796). **B)** Boxplots of expression data from (**A**) showing biological replicates. **C)** DNA methylation averages of indicated cytosine context (y-axis) plotted from transcription start and end sites of TEs that are similarly expressed in *ddm1* and *h1ddm1* (qval >0.05 based on likelihood ratio test (Pimentel et al., 2017) and *ddm1* average expression level >2.5 TPM, n = 284). **D)** Genome browser view of interval showing depletion of DNA methylation at a TE gene promoter and corresponding expression (promoter region highlighted in box). Cytosine context noted in grey. **E)** Methylation averages as in (**C**) of TEs downregulated in *h1ddm1* (qval <0.05 and *ddm1* average expression level >2.5 TPM, n = 210). **F)** Genome browser view of interval showing promoter mCG at a TE gene promoter and corresponding repression of expression in *h1ddm1* but not *ddm1* (promoter region highlighted in box). **G)** Expression level per *ddm1* derepressed TE >2 kb (n=146) and corresponding *h1ddm1* F3 values as a function of their respective promoter mCG, where the promoter region is +/- 500 bp of TSS. **H)** Methylation averages of indicated context (y-axis) plotted from TE TSS (TSS=0 on y-axis; left, categories as from (Osakabe et al., 2021); right, categories determined in this study). **I)** Boxplots of TE expression for published categories (Osakabe et al., 2021, left; neutral: 414, inactive: 381) and categories determined in this study (right, neutral: 450, inactive: 253). **J)** *ddm1* H2A.W (log_2_ ChIP/input) as a function of *ddm1* expression level per TE.

TEs that are similarly expressed between *ddm1* and *h1ddm1* possess a striking, localized depletion of DNA methylation in all contexts near the transcriptional start site (TSS) in *h1ddm1,* so that their methylation levels approach those of *ddm1* plants at the TSS (Fig. 3C-D). In both CG and CHG contexts, average methylation downstream of the TSS increases to an intermediate level within 1 kb (Fig. 3C-D), illustrating that, as in genes (X. Zhang et al., 2006; Zilberman et al., 2007), DNA methylation in the TE body is compatible with high levels of expression. In contrast, TEs with reduced expression in *h1ddm1* compared to *ddm1* show little or no methylation depletion near the TSS and overall non-CG methylation levels similar to *WT* (Fig. 3E-F), indicating that the increased levels of CG and non-CG methylation in *h1ddm1* plants (compared to *ddm1*) attenuate TE expression. Considering *ddm1* and *h1ddm1* together, log_2_ of the expression level shows a strong negative correlation with mCG around the TSS of downregulated TEs (+/-500 bp; TEs longer than 2 kb; *r* = −0.60, Fig. 3G), suggesting that DNA methylation around the TSS quantitatively reduces expression. Taken together, our results support the hypothesis that intermediate mCG gives rise to intermediate TE expression at the cell population level.

### TSS methylation correlates with TE expression in *ddm1* x *WT* F1 plants

Some TEs continue to be highly expressed following *ddm1* outcrossing to *WT* despite DDM1 restoration, whereas others are silenced (Teixeira et al., 2009). A recent study reported that TE silencing in F1 progeny of *ddm1* and *WT* is not correlated with DNA methylation but is instead linked to deposition of the H2A.W histone variant (Osakabe et al., 2021). Analysis of the published TE expression categories indeed shows no methylation difference between silenced (inactive) and expressed (neutral) TEs in the F1 plants (Fig. 3H, left), but these categories also show little difference in TE expression (Fig. 3I and S3D). We therefore calculated expression values from the published raw RNA-seq data to identify 235 TEs that are at least 4-fold downregulated in the F1 plants and compared them to 225 TEs not downregulated in the F1 (Fig. 3H-I and S3E). Consistent with published results (Teixeira et al., 2009), we find a strong recovery of DNA methylation in all contexts around the TSS of re-silenced TEs, but not at expressed TEs (compare “Osakabe et al.” to “reanalysis”, Fig. 3H), as would be expected if TE expression is reduced due to increased DNA methylation.

Considering that loss of DDM1 disperses heterochromatic foci (Soppe et al., 2002), greatly reduces DNA methylation in all contexts (Vongs et al., 1993; Zemach et al., 2013), and strongly activates TE expression (Kato et al., 2004; Miura et al., 2001; Tsukahara et al., 2009), whereas loss of H2A.W does not substantially disperse heterochromatin, alter DNA methylation, or activate TE expression (Bourguet et al., 2021), reduced H2A.W occupancy in heterochromatin is not likely to be a primary cause of the *ddm1* phenotype. Instead, our data indicate that TE activation in *ddm1* mutants is caused by loss of DNA methylation, and reduced TE expression in *h1ddm1* and *ddm1* x *WT* plants is a consequence of increased methylation. As H2A.W occupancy within TE bodies is anticorrelated with TE expression (Fig. 3J), transcription-coupled H2A.W eviction plausibly explains at least some of the observed H2A.W loss from *ddm1* heterochromatin (Osakabe et al., 2021).

### Morphological phenotypes are ameliorated in *h1ddm1* compared to *ddm1*

In addition to extensive disruption of heterochromatin, loss of DDM1 function causes CHG hypermethylation of genes and severe morphological phenotypes (Ito et al., 2015; Kakutani et al., 1995, 1996; Zemach et al., 2013). As heterochromatic methylation and TE silencing are partially restored in *h1ddm1* plants, we evaluated whether other *ddm1* defects are also ameliorated. We find that the widespread *ddm1* genic mCHG increase is largely abolished in *h1ddm1* (Fig. 4A-B). The *ddm1* morphological phenotypes are also markedly improved in *h1ddm1* plants, which have more rosette leaves with more total area, are not delayed in bolting, and grow taller and produce more siliques (fruit) than *ddm1* (Fig. 4C-D). We have propagated *h1ddm1* plants until generation F14, at which point they remain fertile and generally healthy (Fig. S4A). Thus, in addition to stabilized intermediate DNA methylation of heterochromatin, *h1ddm1* plants have a stable, relatively normal morphology in comparison to *ddm1*.

**Figure 4.**
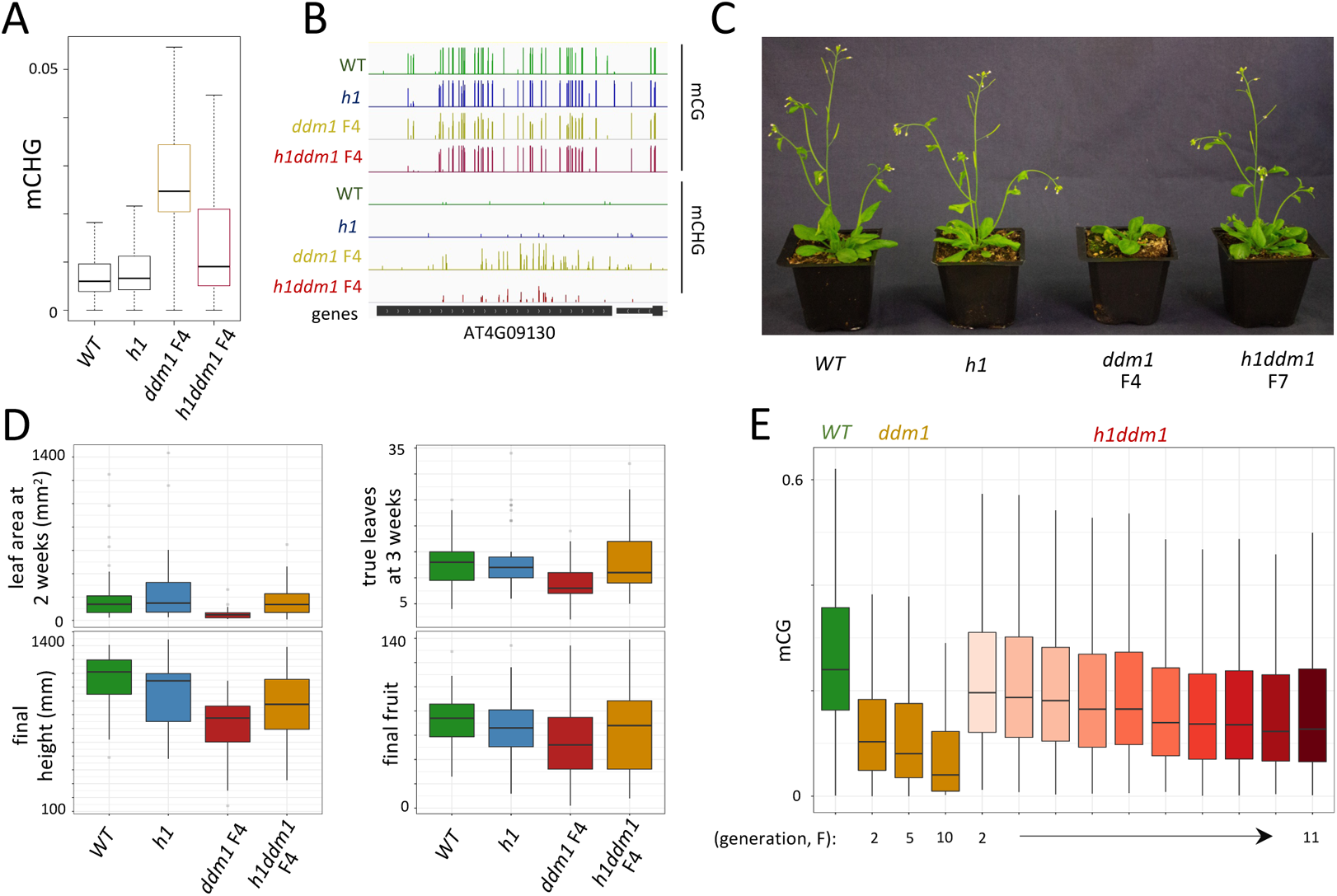
*h1ddm1* rescues diverse *ddm1* phenotypes. **A)** Boxplot illustrating mCHG hypermethylation in *ddm1* (F4) gene-body methylated genes is partially ameliorated in F4 *h1ddm1* (n=561). **B)** Genome browser view of *ddm1* mCHG hypermethylation and its reduction in *h1ddm1* (F4) at AT4G09130. **C)** Plants at 4 weeks post-germination, genotypes are indicated. All genotypes are F4 except *h1ddm1* is F7. **D)** Boxplots of plant measurements conducted on F4 plants of indicated genotypes. Sample numbers per genotype are as follows: *WT*: 43; *h1*: 38; *ddm1*: 35; *h1ddm1*: 49. **E)** Boxplots of mCG in genes hypomethylated in *ddm1* (n=529, p<1.0×10^−13^, Fisher exact test). *ddm1* F10 obtained from (Ito et al., 2015).

### Genic mCG declines across *ddm1* and *h1ddm1* generations

Although DDM1 is primarily important for TE methylation, some genes, especially those with low levels of expression, lose mCG in *ddm1* plants (Fig. S4B) (Ito et al., 2015; Lyons & Zilberman, 2017). Unlike *ddm1* hTEs, in which mCG declines precipitously in the F2 generation and shows little change thereafter (Fig. S1C), these genes show a gradual transgenerational mCG decline (Fig. 4E). As in TEs, mCG in *ddm1*-affected genes is substantially higher in *h1ddm1* than in *ddm1* (Fig. 4E). Compared to TEs (Fig. 1C), genic mCG declines more slowly and stabilizes more gradually across *h1ddm1* generations (Fig. 4E). The slower decline of genic mCG in *ddm1* and *h1ddm1* compared to TEs suggests that mCG maintenance is more robust in genes in the absence of DDM1. This is consistent with most genes retaining roughly *WT* mCG in *ddm1* (Lyons & Zilberman, 2017), so that some loci (hTEs) have strongly compromised mCG maintenance, others (lowly expressed genes) have modestly reduced mCG maintenance, and others (most genes) have effectively normal mCG maintenance.

### Quantification of *h1ddm1* mCG *de novo* and maintenance failure components

The pattern of transgenerational mCG loss and stabilization in *h1ddm1* resembles an exponential decay (Fig. 1C) that should be sensitive to the rates of both maintenance failure and *de novo* methylation. To characterize *h1ddm1* mCG dynamics more precisely, we developed a mathematical model that describes the CG methylation reaction using three processes expressed as linear difference equations: *de novo* methylation per cell cycle (delta, δ), maintenance failure per cell cycle (epsilon, ε), and DNA replication, assuming 34 cell cycles per generation based on published estimates for long day conditions (Fig. 5A and Methods, Watson et al., 2016). Over many cell cycles, any given CG site can be affected multiple times by these processes, dynamically changing back and forth between methylated and unmethylated. Overall, each TE has an initial *WT* mCG level, M_0_, and a lower steady-state mCG level, termed M*, that is approached after multiple generations (*n*) of inbreeding, such that M_n_ → M* for large *n*. The two unknowns δ and ε can be derived on a per-TE basis by fitting the time-series methylation data to an exponential decay, with a timescale [(δ+ε)/2]^−1^, that tends to M*=δ/(δ+ε) (Methods). Longer TEs (>2 kb) proved to be more amenable to modeling due to lower levels of noise. Excluding TEs with an average *WT* mCG ≤ 0.65, the mCG trajectory of the vast majority of TEs >2 kb (1617/1883, 86%) could be reliably fit to an exponential decay (Methods). The fit to the generational averages is shown (Fig. 5B). Although many short TEs were excluded, the remaining TEs comprise ~80% of total heterochromatic sequence length in the *Arabidopsis* genome.

**Figure 5.**
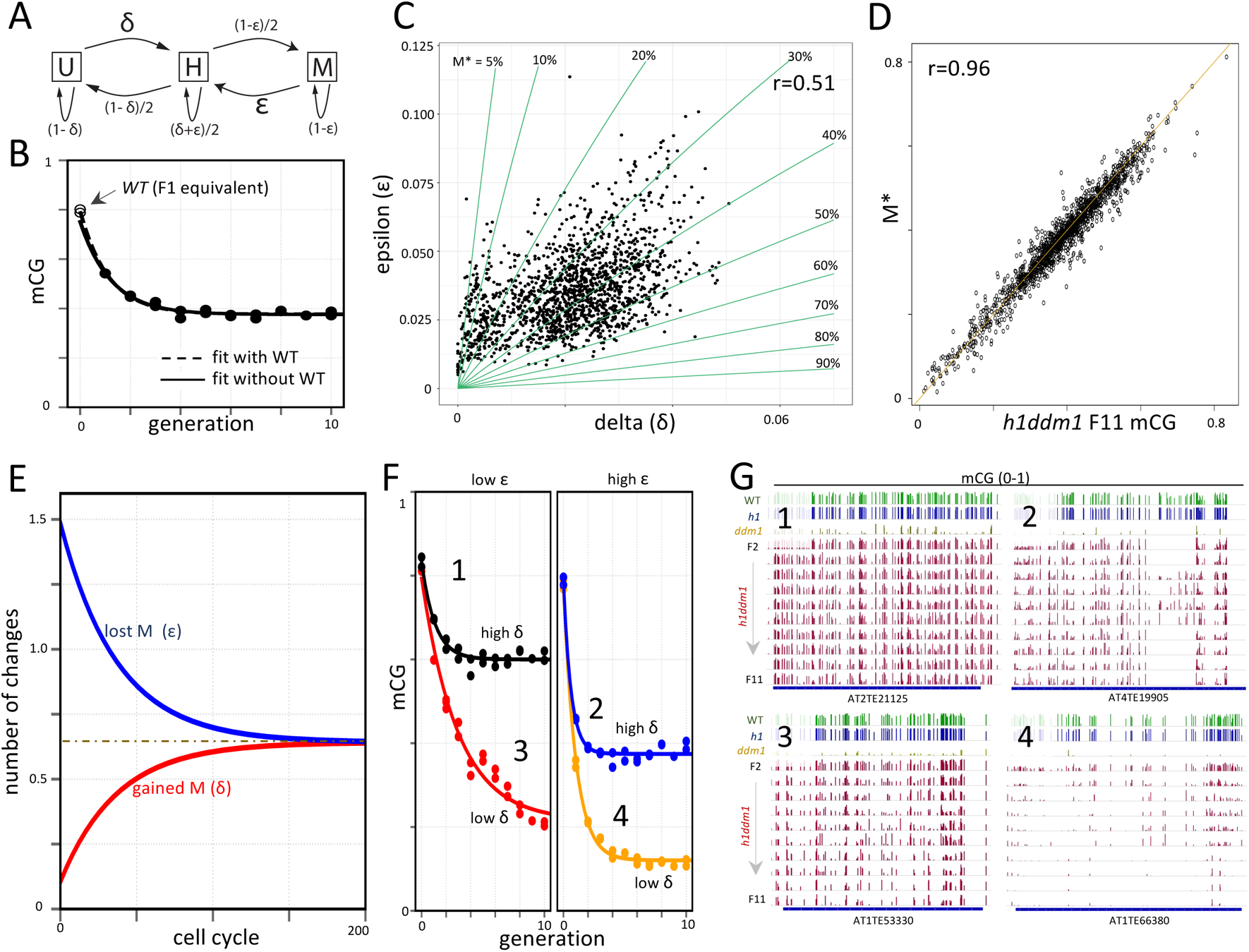
Estimation of per-TE *de novo* and maintenance mCG rates in *h1ddm1*. **A)** Diagram illustrating the relationship between *de novo* methylation (δ), maintenance methylation failure (ε), and the methylation states of a given CG site. U = unmethylated, M = methylated, and H = hemimethylated. **B)** The average mCG of all TEs>2 kb with *WT* mCG>0.65 across *h1ddm1* generations (generation 1 = *h1ddm1* F2) fits an exponential decay. Solid line indicates the fit without including *WT* methylation values as the initial data point (generation 0); dashed line represents the *WT*+*h1ddm1* fit. **C)** Scatterplot of δ vs. ε at those TEs fit by an exponential (n=1617). Green lines indicate the δand ε required to achieve the indicated steady-state methylation level, M*. **D)** Scatterplot of modeled steady-state methylation (M*) vs. *h1ddm1* mCG at F11. **E)** Plot of changes in methylation per cell cycle for a hypothetical TE with 100 CG sites beginning with 90% methylation and with average *h1ddm1* δ and ε values (0.021 and 0.033, respectively). Dotted line illustrates the number of changes at steady-state (0.64). **F)** Different δ and ε regimes produce a variety of mCG dynamics in *h1ddm1*. Quintiles of δ and ε were used to subset the modeled TEs and fit their respective average mCG. Generation 0 = WT, generation 1 = *h1ddm1* F2, etc. **G)** Genome browser views of representative TEs from each of the 4 groups in (**F**). Numbers listed next to curves in (**F**) correspond to browser views shown.

By individually modeling the above TEs, we find that *h1ddm1 de novo* and maintenance failure rates are within comparable ranges and show an intriguing linear correlation of 0.51 (δ_median_=0.022, δ_mean_=0.021, sd=0.010; ε_median_=0.033, ε_mean_=0.035, sd=0.015) (Fig. 5C). We were unable to calculate δ and ε for *ddm1* due to the very rapid loss of mCG in this genotype (Fig. 1A and S1C). However, we find that TEs that most readily regain mCG in *ddm1* x *WT* F1 plants (Osakabe et al., 2021) have the highest δ values in *h1ddm1* (Fig. S5A, r=0.48), suggesting that the *h1ddm1* δ values are relevant for other genotypes.

By altering the values of δ and ε, essentially any steady-state methylation level can be achieved, as shown in the breadth of observed M* at individual TEs, ranging from near 0 to greater than 60% (Fig. 5D). Further, different δ and ε combinations can achieve the same M*. To help visualize this, we overlaid lines on a plot of ε vs. δ that correspond to given steady-state mCG states (M*) (green lines, Fig. 5C). These lines emphasize that a low ε (efficient maintenance) is usually required for the *WT* mCG level, and that even a small increase of the absolute maintenance failure rate can have a substantial impact on the steady-state methylation level. Thus, although the average *h1ddm1* ε is only about 3%, this compounds over many cell cycles to produce a substantially lower steady-state mCG compared to *WT* (Fig. 1A-C).

To appreciate the implications of this model, consider a TE comprised of 100 CG sites, beginning with 90% methylation, and where δ and ε are equal to their *h1ddm1* mean values. *De novo* activity affects unmethylated CG sites (uCG), and therefore plays only a small role in the initial cell cycles following the creation of the mutant state (generation 0, only 10 uCG, Fig. 5E). However, as time progresses, unmethylated sites increase in prevalence and the absolute contribution of *de novo* methylation increases, whereas maintenance failure becomes relatively less important as the number of mCG sites decreases. A steady-state is approached around cell cycle 150 (corresponding to just over four generations), when the numbers of gains and losses are balanced, both converging to a constant equal to (on average) 0.64 CG sites gaining/losing per cell cycle (dashed line in Fig. 5E).

Different regimes of δ and ε manifest as different steady-state levels and/or mCG trajectories (with faster or slower decays) (Fig. 5F-G). Efficient maintenance and low *de novo* each cause a slower decay to the mCG steady-state, so that TEs with excellent maintenance and low *de novo* fail to reach steady-state even after 10 generations (Fig. 5F-G, #3, n=146 TEs), whereas those with highest *de novo* but poor maintenance converge to steady-state rapidly (Fig. 5F-G, #2, n=149 TEs). At the most mCG-depleted TEs, *de novo* rates are low, with larger maintenance failure, and average mCG stabilizes at ~10% (Fig. 5F-G, #4, n=17 TEs). At the other end of the spectrum, TEs with high δ and low ε lose ~20% of their *WT* mCG (Fig. 5F-G, #1, n=13 TEs). However, because δ and ε are positively correlated (Fig. 5C), TEs with poor maintenance tend to have better *de novo*, and vice versa, so that the above extremes (Fig. 5F-G, #1 and #4) are unusual. Although we could not fit genic methylation dynamics well enough to confidently quantify δ and ε, the more gradual decline of mCG in genes (Fig. 4E) than in TEs (Fig. 1C) is consistent with lower δ and ε (less *de novo* activity and better maintenance) in genes.

### Contributions of DRM and CMT methyltransferases to CG methylation in *h1ddm1*

Our model indicates that the initial mCG decline is dominated by maintenance failure, whereas steady-state mCG has a strong dependence on *de novo* methylation (Fig. 5E-F). The correlation of late generation (steady-state) mCG with mCHG and mCHH (Fig. 1E) therefore suggests that the RdDM and/or CMT pathways that mediate mCHG/CHH contribute (directly or indirectly) to *de novo* mCG activity. Consistent with this, δ shows strong linear correlations with mCHG and mCHH (r_mCHG_=0.66, r_mCHH_=0.59) (Fig. S5B-C). To elucidate the enzymatic origins of *de novo* mCG, we generated three *h1ddm1* mutant lines, each lacking one of the three principal *Arabidopsis* non-CG methyltransferases: DRM2, CMT2 and CMT3. We inbred each line through the F7 generation and analyzed leaf methylomes as we did for *h1ddm1*. The *h1ddm1drm2* and *h1ddm1cmt2* lines were created in the same way as the initial *h1ddm1* line: a plant homozygous for all mutations except *ddm1* was allowed to self-fertilize to create the founder F2 in which the *ddm1* mutation was homozygous for the first time. We were unable to generate *h1ddm1cmt3* plants this way, and instead created an F2 *h1ddm1*(*cmt3*/+) plant (first generation of *ddm1* homozygosity), which was allowed to self-fertilize to create F3-equivalent *h1ddm1cmt3* plants (first generation of *cmt3* homozygosity).

Average hTE mCG is greatly reduced in *h1ddm1drm2* compared to *h1ddm1*, which becomes obvious after the F2 generation (Fig. 6A-B). Consistently, the overall morphology of *h1ddm1drm2* plants resembles *ddm1* more than *h1ddm1* (Fig. S6A). In contrast, neither *h1ddm1cmt2* nor *h1ddm1cmt3* lines exhibit an overall mCG decrease compared to *h1ddm1* (Fig. 6A-B), despite corresponding strong losses of mCHH and mCHG, respectively (Fig. S6B-C), and these lines morphologically resemble *h1ddm1* (Fig. S6A).

**Figure 6.**
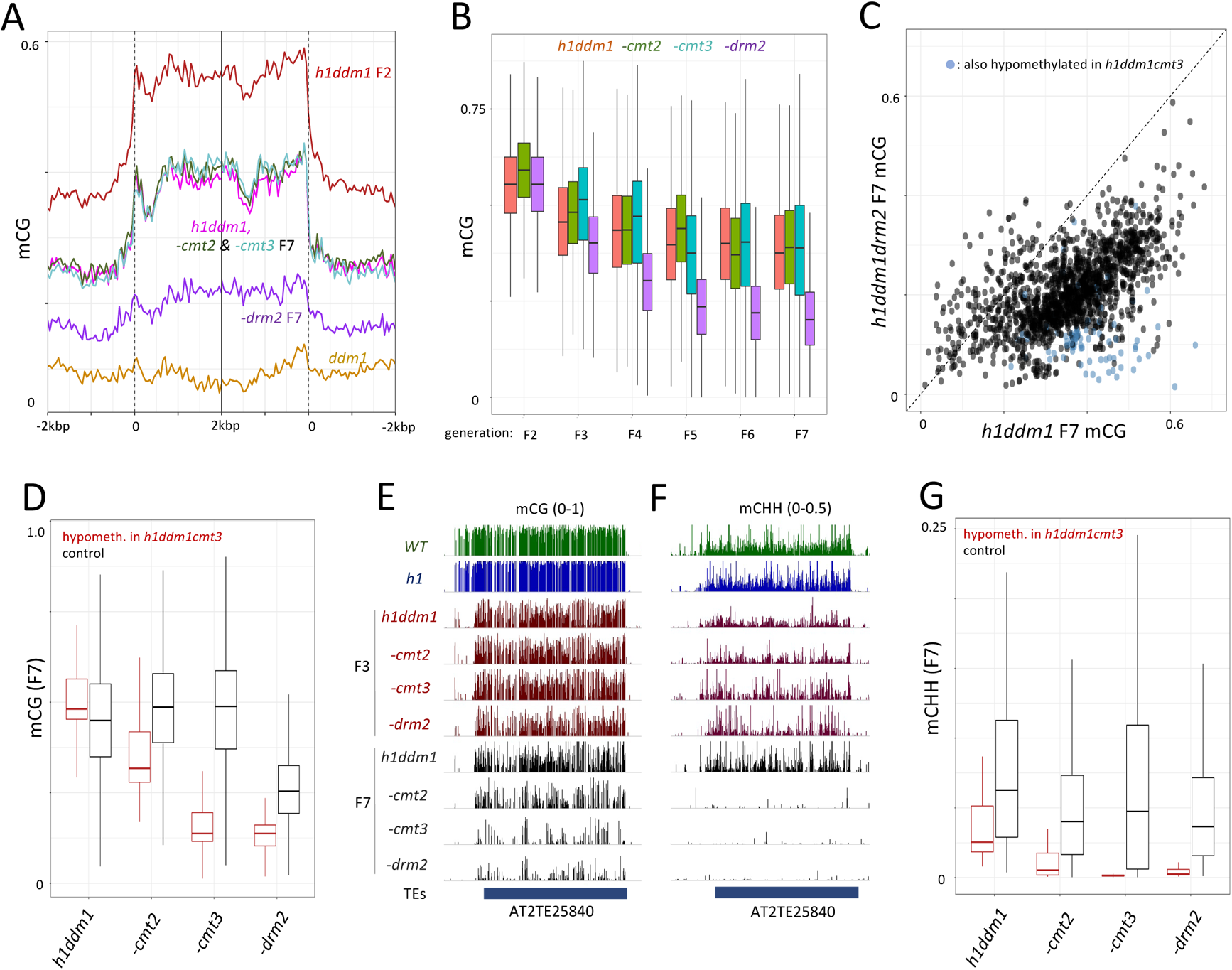
Steady-state mCG in *h1ddm1* depends on DRM2. **A)** Average mCG is plotted for all heterochromatic TEs for indicated genotypes around TE start and stop sites (TEs >250 bp). *h1ddm1*, *h1ddm1cmt2*, and *h1ddm1cmt3* resemble one another at F7, whereas mCG is much reduced in *h1ddm1drm2*. **B)** Boxplots of per-TE mCG over seven generations for *h1ddm1* and compound *h1ddm1* methyltransferase mutants (*h1ddm1cmt3* F2 is absent; please see text). **C)** Scatterplot comparing *h1ddm1drm2* F7 to *h1ddm1* F7 average mCG of previously modeled TEs (n=1619), with blue points highlighting the TEs significantly hypomethylated in *h1ddm1cmt3* relative to *h1ddm1* (n=86, p-val <0.01, Fisher exact test). **D)** Boxplots comparing mCG in modeled TEs that are either hypomethylated in *h1ddm1cmt3* (red) or control TEs that are not significantly hypomethylated (black). **E-F**) Genome browser view illustrating mCG (**E**) and mCHH (**F**) at TE with CMT3-dependent mCG. **G**) Boxplots comparing mCHH levels as in (**D**). Note the near complete depletion of mCHH in *h1ddm1cmt3* but not -*cmt2* at CMT3-depdnent TEs, indicating CMT3 is required for DRM2 activity at these loci.

Although the loss of CMT3 does not substantially reduce average heterochromatic mCG, we identified 86 TEs with mCG in *h1ddm1cmt3* mutants comparable to *h1ddm1drm2* (>50% mCG decrease; p < 0.01, Fisher’s exact test, Fig. 6C-E and S6D). Because non-CG methylation promotes RdDM (Choi et al., 2021; Johnson et al., 2014; Liu et al., 2014), we suspected the loss of CMT3 might reduce RdDM activity at these loci. Indeed, mCHH (the hallmark of RdDM) is nearly eliminated at these TEs in F7 *h1ddm1cmt3* plants, as it is in F7 *h1ddm1drm2* (Fig. 6F-G, mCHG shown in Fig. S6E), indicating that RdDM activity depends on CMT3 at a subset of heterochromatin. These CMT3-dependent TEs also have reduced mCHH in F7 *h1ddm1cmt2* plants (Fig. 6F-G), as well as reduced mCG (Fig. 6D-E), so that steady-state mCG is affected by all three of the tested methyltransferases. Notably, non-CG methylation at these TEs does not collapse immediately in the compound methyltransferase mutants, but can be similar to *h1ddm1* in early generations, and tends to decrease in tandem with mCG (Fig. 6F and S6F-H). This is consistent with our observation that CG and non-CG methylation changes are correlated across *h1ddm1* generations (Fig. 1E), and with the known dependence of non-CG methylation on mCG at some TEs (Choi et al., 2020; Choi et al., 2021; Y. Zhang et al., 2018). The CG and non-CG methylation decreases may therefore be mutually reinforcing. Together, these data demonstrate that DRM2 is a key contributor to intermediate mCG in *h1ddm1* heterochromatin, but also that epigenetic inheritance of mCG involves integration of the RdDM, CMT and MET1 pathways, at least at some loci.

### DRM2 mediates most of the *de novo* mCG in heterochromatin

Application of our discrete methylation model to the methyltransferase mutant data allows us to calculate the relative contributions of each methyltransferase to *de novo* and maintenance mCG (Fig. 7A-C). Surprisingly, we find that ε decreases for all three compound mutants compared to *h1ddm1* (Fig. 7D), indicating maintenance becomes more efficient when these methyltransferases are removed. This is likely related to the observation that F2 *h1ddm1* mCG shows a strong negative correlation with H3K9me2 (Fig. 1D). As the initial mCG decline is dependent in large part on ε (Fig. 5E-F), ε should also correlate with H3K9me2, which is indeed the case (Fig. 7E). Because non-CG methylation and H3K9me2 vary in tandem (Stroud et al., 2014), our data suggest that the *drm2*, *cmt2* and *cmt3* mutations improve mCG maintenance by rendering TEs less heterochromatic (less non-CG methylation and H3K9me2) and therefore less dependent on DDM1.

**Figure 7.**
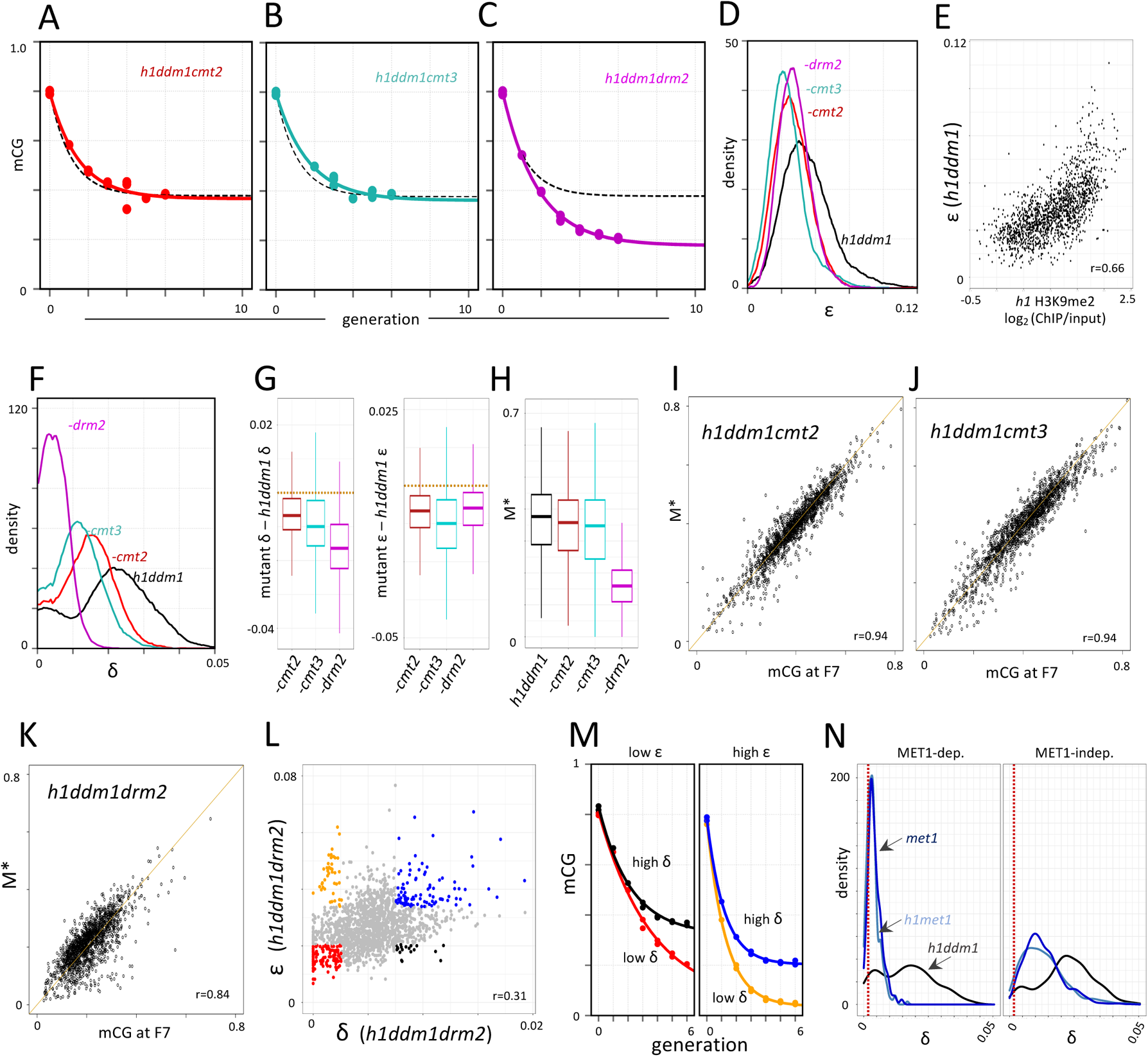
Estimation of mCG rates in compound *h1ddm1* methyltransferase and *met1* mutants. **A-C**) Comparison of average modeled TE mCG decay fits (colored by genotype) to *h1ddm1* (black dashed lines). **D)** Density plots of modeled ε for individual TEs. **E)** Scatterplot showing *h1ddm1* ε as a function of parental *h1* H3K9me2. **F)** Density plots of modeled δ (see Methods regarding the low-δ peaks). **G)** Boxplots of the change in δ (left) or ε (right) for the indicated genotypes. Orange horizontal lines are at y=0. **H)** Boxplots of modeled steady-state methylation (M*) for the indicated mutants. **I-K**) Scatterplots of modeled steady-state methylation (M*) vs. mCG at F7 for the indicated genotypes, with Pearson’s r shown. **L)** Scatterplot of δ vs. ε in *h1ddm1drm2* TEs, with differently colored points indicating the different extremes possible in this mutant, grouped per quintile combination. **M)** Color-matched exponential decay fits to average mCG of the quintile combinations shown in (**L**). Generation 0 = *WT*, generation 1 = *h1ddm1drm2* F2, etc. **N)** Density plots depicting the distributions of δ for the indicated genotypes in either MET1-dependent (left, n=271) or -independent (right, n=1258) modeled TEs (see Methods). The 5.5% of modeled TEs not in these categories were excluded from this analysis. Red dashed lines indicates the estimated false-positive methylation rate for the *met1* and *h1met1* data (see text).

The *cmt* compound mutants exhibit modest decreases in δ that are comparable to the decrease in ε. In *h1ddm1cmt2,* δ decreases by 33% (from 0.021 to 0.014) and ε decreases by 26% (from 0.035 to 0.026), and in *h1ddm1cmt3,* δ decreases by 43% (to 0.012) and ε decreases by 34% (to 0.023; Fig. 7D,F-G). The changes between δ and ε are correlated (Fig. S7A-C), resulting in a form of dynamic compensation that, in the case of *cmt* mutants, keeps steady-state mCG (which depends on the ratio of δ and ε) nearly the same as in *h1ddm1* (Fig. 7H-J). However, in *h1ddm1drm2*, δ falls 75% (to 0.0053; Fig. 7F-G) but maintenance does not improve at a similar level (ε decreases by 23% to 0.027; Fig. 7D,G). The much greater relative decrease in δ causes greatly reduced steady-state mCG compared to *h1ddm1* (Fig. 7H,K) and accordingly, TEs with both high δ and low ε are very uncommon in *h1ddm1drm2* as compared to low δ and high ε (Fig. 7L-M, black and orange points, respectively). Our results indicate that although CMT2, CMT3 and DRM2 all participate in the epigenetic dynamics of heterochromatic mCG, DRM2 contributes the bulk of the *de novo* mCG activity.

### *De novo* rate estimates are consistent between *h1ddm1* and *met1*

Because we calculate TE *de novo* mCG rates in the *h1ddm1* background, it is important to ascertain how relevant these rates are for *WT*, especially because loss of histone H1 substantially perturbs RdDM activity (Choi et al., 2021; Papareddy et al., 2020). To address this issue, we calculated *de novo* rates in *met1* and *h1met1* mutants using published data (Choi et al., 2020). These mutants are assumed to lack maintenance activity (ε=1), and δ values can therefore be straightforwardly inferred based on measured steady-state mCG (δ=mCG, Methods). The *met1* and *h1met1* δ values are correlated with *h1ddm1* δ values (Fig. S7B-C, r=0.28 for *met1* and r=0.34 for *h1met1*), supporting the conclusion that *h1ddm1* δ values are relevant for other genotypes. Some TEs retain RdDM and CMT activity in *met1* and *h1met1* mutants (MET1-independent TEs), whereas others lose methylation in all sequence contexts (MET1-dependent TEs; Choi et al., 2020; Choi et al., 2021; Y. Zhang et al., 2018). As expected, *h1ddm1* δ values show similar distributions at these TE categories (Fig. 7N). However, the *met1* and *h1met1* δ values are much higher in MET1-independent TEs, where they are much more comparable to those for *h1ddm1* (Fig. 7N), as would be expected if *de novo* activity is mediated by the non-CG methylation pathways.

Bisulfite sequencing has an intrinsic error rate caused by incomplete chemical conversion of unmethylated cytosine, as well as PCR and sequencing errors (Krueger & Franke, 2012). This rate can be estimated based on the measured methylation frequency in the unmethylated chloroplast genome, and is 0.17% for both *met1* and *h1met1* (dashed lines in Fig. 7N). Because per-TE error rates will be distributed around 0.17%, *met1* and *h1met1* δ estimates that are close to this value should be treated as upper bounds, with real δ values potentially much smaller. Therefore, many MET1-dependent TEs may experience very low *de novo* rates in *met1* and *h1met1* plants (Fig. 7N, left panel). In contrast, MET1-independent TEs generally have δ values well above the error rate (Fig. 7N, right panel), with the average *met1* and *h1met1* δ values (0.013 for both) in MET1-independent TEs only 1.7-fold lower than that for *h1ddm1* in MET1-independent TEs (δ_mean_=0.022). Because RdDM activity at MET1-independent TEs is similar between *met1* and *WT* (Choi et al., 2021), *WT* heterochromatic *de novo* rates should be in a comparable range.

## Discussion

Despite the similarities between the reported rates of mCG change in *C. neoformans* (Catania et al., 2020) and *Arabidopsis* (Becker et al., 2011; Hazarika et al., 2022; Schmitz et al., 2011; Van Der Graaf et al., 2015), our data indicate that epigenetic inheritance of mCG in *Arabidopsis* TEs involves a strong *de novo* component comparable to mammalian systems (Haggerty et al., 2021; Lövkvist et al., 2016; Wang et al., 2020). The *de novo* activity is primarily mediated by RdDM, with smaller contributions from CMT2/3. Both of these pathways have self-reinforcing features that focus their activities on TEs (Choi et al., 2021; Du et al., 2014; Johnson et al., 2014; Liu et al., 2014; Rajakumara et al., 2011), thereby solving the problem of targeting *de novo* mCG to regions with existing methylation. We derive our *de novo* rates primarily from *h1ddm1* data, but they are in good agreement with the rates in *met1* mutants (Fig. 7N), and should therefore be a good estimate for *WT de novo* rates.

The *de novo* mCG rates we find in heterochromatin are about 1% per site per cell cycle (δ/2, Fig. 5, Methods), which translates to around 30% per generation (1 - 0.99^34^). These rates are at least three orders of magnitude higher than the overall *de novo* rate of 3.2×10^−4^ per generation reported for TEs (Van Der Graaf et al., 2015) and the range of 2.3×10^−6^ to 1.7×10^−4^ recently reported within different TE-associated chromatin states (CSs; Hazarika et al., 2022). This large difference is likely accounted for by two related factors. First, CS rates were calculated using the AlphaBeta model (Shahryary et al., 2020) that produces a good fit for CSs associated with genes, explaining up to 88.6% of the data variance in CS5 (Hazarika et al., 2022). However, the model performs less well with the TE-associated states CS30-36. AlphaBeta provides reasonable fits for CS30 (67.6% of variance explained) and CS31 (57% of variance explained), and these states have relatively high reported *de novo* rates (3.5×10^−5^ for CS30 and 1.7×10^−4^ for CS31). AlphaBeta does not handle other TE CSs well, ranging from 28.3% of variance explained for CS32 to effectively 0% of variance explained for CS36 (variance explained values are from Supplementary Table 2 of Hazarika et al.). This indicates that the AlphaBeta model is not reliable for these regions of the genome.

More generally, AlphaBeta and other published rate estimates rely on the assumption that the rates are low enough that all or nearly all changes are captured by measurements that are separated by one or several generations. This should be a good assumption for genes, but it is not for TEs. Per-generation *de novo* rates of 30% mean that the half-life of an unmethylated site is short (30% rate implies a half-life of about two generations), leading to underestimates of methylation failure rates, because CG sites that lose methylation are rapidly remethylated. This likely at least partially explains why the reported per-generation maintenance failure rate for genes (1.5×10^−3^) is 100-fold higher than the rate for TEs (1.2×10^−5^; Van Der Graaf et al., 2015). Calculation of *de novo* rates may be confounded by an even more serious issue. Because *de novo* rate calculations rely on ancestrally unmethylated CG sites, and the half-life of unmethylated sites is inversely related to the *de novo* rate, CG sites with unusually low *de novo* rates will dominate the calculation, leading to potentially very large rate underestimates. This might explain the 1000-fold difference between the reported per-generation *de novo* rate in TEs (3.2×10^−4^; Van Der Graaf et al., 2015) and our estimate of around 3×10^−1^.

Our results indicate that the low levels of mCG divergence observed at plant TEs (Hazarika et al., 2022; Van Der Graaf et al., 2015) are not caused by especially low rates of methylation loss and gain. Instead, this phenomenon is caused by a very high *de novo* rate that is much greater than the maintenance failure rate, so that maintenance errors are rapidly corrected – an idea consistent with the earlier proposal that TE mCG is stabilized by an epigenetic process that favors methylation gain over methylation loss (Van Der Graaf et al., 2015). Our data indicate that stable epigenetic inheritance of TE mCG is ensured by a combination of DDM1-supported MET1 maintenance activity with the strong *de novo* activity of RdDM (with smaller contributions from the CMT pathway). Using the reported per-generation maintenance failure rate for genes (1.5×10^−3^) as a baseline, the *de novo* activity at TEs is strong enough to stabilize mCG in *h1ddm1* despite a >100-fold increase in the maintenance failure rate, to ~4×10^−1^ per generation (equivalent to 1 - 0.985^34^, Fig. 5, per-cell cycle loss rate=ε/2, Methods). Importantly, there is no compelling reason to assume that maintenance efficiency is the same or similar between genes and TEs in *WT* plants. Maintenance efficiency at TEs could be higher than at genes, but – even with the support of DDM1 – may also be considerably lower, especially at heterochromatic TEs. One of the main conclusions of our study is that the high *de novo* rates in TEs enable stable epigenetic inheritance of mCG within a wide range of maintenance rates.

Involvement of the RdDM and CMT pathways implies that *de novo* mCG rates likely fluctuate during development, as substantial differences in the activities of these pathways have been described across cell types and tissues (Erdmann & Picard, 2020), including during *Arabidopsis* reproductive and embryonic development (Bouyer et al., 2017; Calarco et al., 2012; Ibarra et al., 2012; Kawakatsu et al., 2017; Long et al., 2021; Zhou et al., 2022). Embryonic development is characterized by a gradual rise of mCHH in TEs that peaks at the onset of germination and is mediated by RdDM and CMT2 (Bouyer et al., 2017; Kawakatsu et al., 2017). The functional significance of this process has been unclear, and one of its consequences may be to reinforce mCG inheritance by restoring CG sites within TEs to a methylated state.

Our results also bear on the functionality of DDM1 and of plant heterochromatin more broadly. The remarkable alleviation of the *ddm1* phenotype (TE activation, decreased fecundity, increased gene body mCHG, etc.; Fig. 3A-F and 4A-D) that emerges in *h1ddm1* concomitantly with the restoration of heterochromatic DNA methylation illustrates the central importance of chromatin homeostasis in the regulation of organismal viability, and the crucial role of DNA methylation in heterochromatin homeostasis and function. This is consistent with the reports that the mere presence of sufficiently large segments of demethylated heterochromatin in genetically *WT* plants is sufficient for phenotypic disruption (Johannes et al., 2009; Kooke et al., 2015). Our data do not support the proposal that the *ddm1* phenotype, including TE activation, is primarily caused by loss of the H2A.W histone variant from heterochromatin (Osakabe et al., 2021) – a conclusion that is also inconsistent with the published phenotypes of *ddm1* and *h2a.w* loss-of-function mutants (Bourguet et al., 2021; Jeddeloh et al., 1999; Kakutani et al., 1995, 1996; Tsukahara et al., 2009; Zemach et al., 2013). Instead, our results indicate that *ddm1* phenotypes are primarily caused by loss of heterochromatic DNA methylation, and the ameliorative effects of H1 removal in the *ddm1* background stem directly from increased heterochromatic DNA methylation, especially steady-state mCG.

## Materials and Methods

### Biological materials and growth conditions

The *h1ddm1* line was previously described (Lyons & Zilberman, 2017). DDM1 is a potent regulator of methylation and its removal initiates the decay of mCG in *h1ddm1*. Therefore, to generate *h1ddm1* we self-fertilized *h1/+;ddm1/+* and isolated *h1;ddm1.* To generate compound *h1ddm1* methyltransferase mutants, we crossed *h1/h1;ddm1*/+ (abbreviated *h1;ddm1/+*) to homozygous null *drm2-2* (SALK_150863C)*, cmt2-3* (SALK_012874) and *cmt3-11* (SALK_148381C) mutants. We then isolated *h1;drm2;ddm1/+* and *h1;cmt2;ddm1/+* and self-fertilized these to generate each respective F2 *drm2* and *cmt2* compound mutant. For *cmt3* we were unable to obtain *h1;cmt3;ddm1/+* from the initial cross. We did find *h1;ddm1;cmt3/+* frequently however, and therefore used this to generate *h1ddm1cmt3,* but as a result were unable to generate an equivalent F2 for this line. *Arabidopsis* seeds were sown on soil, stratified for 4 days at 4 deg. C and transferred to controlled environment chambers where they were grown in 16h light / 8hr dark at 20 deg. C until tissue harvest.

### Leaf DNA isolation and bisulfite library prep

DNA was isolated by pulverizing ~0.5 g of flash frozen rosette leaves of 4 week post-germination plants. ~100 mg of resulting powder was used to extract genomic DNA (gDNA) with the DNeasy plant mini kit (Qiagen cat. no. 69104) per manufacturer’s instructions. gDNA was subsequently sonicated with the Bioruptor Pico (Diagenode) to ~250 bp median fragment length using 10 cycles of 30 seconds on and off. Agencourt Ampure beads (referred to as “beads” henceforth, cat. no. A63881) were then used at 2X volume to purify the sheared DNA. Following ligation of methylated Truseq sequencing adapters (Illumina) to sheared DNA, bisulfite conversion of DNA was carried out according to manufacturer’s protocol (Qiagen Epitect Kit, cat. no. 59104) except without using carrier RNA. DNA was purified twice with 1.2X beads and converted a second time to ensure complete bisulfite conversion of unmethylated cytosine. Libraries were constructed using NEBnext kits (NEB cat. no. E7645) or Nugen/Tecan Ovation Ultralow (cat. no. 0344NB-08) following the manufacturer’s instructions. NEB next indexing primers (cat. no. E7335S) were used for generating multiplexed libraries during PCR amplification.

### Sperm DNA isolation and bisulfite library prep

Open flowers of F4 or F5 plants (which is here denoted as F4 or F5 sperm, respectively) were collected for pollen isolation in Galbraith buffer (45 mM MgCl2, 30 mM sodium citrate, 20 mM MOPS, 1% Triton-X-100, pH7.0) by vortexing at 2000 rpm for 3 min. Flower parts were removed by straining through a 40 micron filter. Pollen grains were obtained by centrifugation at 2600 g for 5 minutes and broken down with glass beads (Sigma). The lysate was then transferred to a 40 micron cell strainer and the flow-through was centrifuged at 800 g for 10 min at 4°C. The pellet was re-suspended in Galbraith buffer and stained with SYBR Green for fluorescence-activated cell sorting (FACS). Vegetative nuclei and sperm nuclei (SN) were separated and collected based on size and fluorescence intensity. DNA was extracted from SN with ChargeSwitch gDNA Micro Tissue Kit (ThermoFisher cat. no. CS11203) following the manufacturer’s instructions. Briefly, SN were lysed in lysis buffer with proteinase K provided by the kit in a 55°C water bath overnight. RNA was digested with RNase A at room temperature for 30 minutes, then DNA was captured with magnetic beads provided, washed, and eluted. After quantification by fluorometry, about 10-20 ng of DNA was used for bisulfite-sequencing library preparation with Ovation Ultralow Methyl-Seq DR Multiplex System (Nugen part no. 0336) following the manufacturer’s instructions. Briefly, DNA was sonicated with Bioruptor sonicator, end repaired, ligated to adapters which contain indexes, and bisulfite converted twice with EpiTect Fast DNA Bisulfite Kit (Qiagen 59802) following the manufacturer’s instructions. After PCR amplification, libraries were purified with Agencourt RNAClean XP Beads.

### Leaf RNA isolation and library preparation

Leaves (as above) from the F3 generation were flash frozen in liquid nitrogen, pulverized with mortar and pestle on dry ice, and the resulting material was subjected to vortexing in Trizol (Invitrogen, cat. no. 15596-026). Chloroform was then added at one-fifth the total volume and further vortexing was carried out until the solution appeared homogenous. RNA was subsequently pelleted in ice-cold isopropanol. The resuspended RNA was subjected to rRNA removal with Ribo-zero plant kit (Illumina, MRZPL1224) according to the manufacturer’s protocol. 50ng ribo-depleted RNA was used for library preparation with the Scriptseq kit (Epicentre, cat. no. SSV21124) following the manufacturer’s protocol but with the following modifications: the RNA fragmentation step was extended to 10 minutes, and the temperature was increased to 90°C.

### Sequencing

For *h1ddm1* bisulfite libraries and some control and compound *h1ddm1* methyltransferase libraries, we used the Illumina HiSeq 2500 at the QB3 Vincent Coates Genomic Sequencing Lab at UC Berkeley. *h1ddm1* compound mutants were mostly sequenced on the NextSeq 500 (Illumina) at the John Innes Centre. Sperm bisulfite libraries were sequenced at the Bauer Core Facility at Harvard University with Illumina HiSeq 2000.

### Short read mapping and quantification

Bisulfite libraries were mapped to the genome with BSMAP (Xi & Li, 2009) for all analyses, except single-read analyses, for which we used Bismark (Krueger & Andrews, 2011). BSMAP output was converted to per-base methylation scores with BSMAP’s methratio.py script. RNA was mapped to the genome with Hisat2 (D. Kim et al., 2019), using default settings except “dta” was on. Transcripts were assembled using Stringtie (Pertea et al., 2015) using Araport11 (March 2021 release; available as Araport11_GTF_genes_transposons.Mar202021.gtf at www.aradibopsis.org) as a guide, as we noticed that *ddm1* mutant transcripts added in this release (Panda & Slotkin, 2020) were also present in our *ddm1* data. For subsequent RNA quantification, we used our merged assembled transcripts to generate indices for use with kallisto pseudomapping (Pimentel et al., 2017) using the following settings for the quant program: --single --fr-stranded -b 100 -l 320 -s 30.

### Statistics and visualization of experimental data

Methratio.py output was converted to GFF and further processed and analyzed using commands and workflows outlined in https://github.com/dblyons/modeling_h1ddm1/.

Kallisto output was fed into the R environment for processing and fitting with the Sleuth software package, which was used to perform likelihood ratio tests to derive q-values (i.e. FDR-adjusted p-values) for each TE per genotype comparison, as outlined here: https://rawgit.com/pachterlab/sleuth/master/inst/doc/intro.html.

The ends-analysis.pl script (https://zilbermanlab.net/dzlab-tools-1-5-81-linux-tar/) was used to generate enrichment score matrices of mapped bisulfite data around genomic features of interest. These matrices were imported to R (http://www.R-project.org) for further processing and visualization using code available at www.github.com/dblyons/R. Both base functions and the ggplot2 library (Wickham 2009) were used for all plots except for heatmaps in Fig. 1, Fig. 2 and Fig. S1, and genome browser images. Heatmaps were generated with pheatmap (https://cran.r-project.org/web/packages/pheatmap/index.html). Built-in R function ‘cor’ was used for calculating Pearson’s correlation. Genome tracks are screenshots of our data were displayed in Integrated Genome Viewer (Robinson et al., 2011). For RNA-seq, we used bedtools genomecov (Quinlan & Hall, 2010) with scaling on to convert Hisat2 bam output to scaled browser tracks with units in reads per million mapped (RPM). For determining MET1-dependency of mCHH, we used the same method as in (Choi et al., 2021) such that: TE mCHH > =5% in WT and > =5% in *met1* is MET1-independent, while TEs fitting the following criteria are MET1-dependent: TE mCHH > =5% in WT and < 2% in *met1*.

### Plant leaf area measurements

Flats of the four different genotypes of plants were arranged on a platform to align images. Each plant was segmented manually using ImageJ (Collins, 2007), and then analyzed using ImageJ’s mask feature to isolate leaves; the measure feature was used to measure all visible leaf area. Photos were taken using a Canon Rebel T3i.

### Reanalysis of data from (Osakabe et al., 2021)

To determine whether DNA methylation changes occur at the promoters of TE genes that are significantly repressed in the *ddm1* x Col-0 hybrid, we used the expression values provided in source data for Figure 5 (https://www.nature.com/articles/s41556-021-00658-1#Sec36, Osakabe et al., 2021) to group TE genes as *ddm1* active, and if active, downregulated (inactive) or similarly expressed (neutral) in the *ddm1* x Col-0 hybrid F1. We then performed new transcript abundance analysis using kallisto and sleuth (as described above) without using any clustering as was done in (Osakabe et al., 2021), and then generated TE groupings based on the ratio of hybrid/*ddm1* TPM as follows (as shown in Fig. S3E): downregulated < 0.25 and q-val < 0.05 (n=253); neutral > 0.3 (n=450). For mapping of methylation at these groups of TEs, we calculated the difference of observed methylation in the hybrid minus the mean of the parental lineages (Col-0 + *ddm1*) / 2 as 50 bp bins across the relevant loci, and then doubled this number to account for the dilution of the sample by the *WT* alleles present in the sample. We used the same calculation for estimating mCG gained at the *ddm1* allele in Fig. S5A, except for the doubling step. For comparison of *ddm1* H2A.W with TE expression, we used H2A.W ChIP-seq processed data from GSE150436 and plotted these against our recalculation of *ddm1* TE gene expression, which was remarkably similar to the published results (Osakabe et al., 2021). For the calculation of gained mCG in the hybrid shown in Fig. S5A, we used the difference of observed methylation in the hybrid minus the mean of the parental lineages (Col-0 and *ddm1*) from (Osakabe et al., 2021).

### Transposable element annotations

Heterochromatic TEs used for methylome analysis here are the same as in (Choi et al., 2020). For *ddm1* mutant transcriptional analysis shown in Figure 3, we use the assembled transcripts from our analysis outlined above, which closely matches that of the March 2021 Araport11 gene and transposon release. These annotations, as well as genome coordinate files of the modeled TEs, TEs upregulated in *ddm1*, and the Osakabe et al. reanalysis annotations are available at www.github.com/dblyons/annotations_2022.

### Mathematical model for TE methylation dynamics: constructing the recursion relations

We model the transgenerational methylation dynamics of CG sites within TEs. Each CG site is defined to be in one of three states: fully unmethylated, hemi-methylated or fully methylated. Recursion relations are used to express the methylation status of the TE at the end of each cell cycle such that:

*U^(n)^* = fraction of sites unmethylated at the end of the *n^th^* cell cycle

*H^(n)^* = fraction of sites hemi-methylated at the end of the *n^th^* cell cycle

*M^(n)^* = fraction of sites methylated at the end of the *n^th^* cell cycle

where *U^(n)^* + *H^(n)^* + *M^(n)^* = 1.

As each CG site contains two cytosines, our model relates to the experimentally measured methylation level, 〈*mC*〉, via 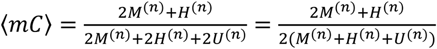.

To derive the recursion relations, we consider three key processes during the cell cycle: a) replication, b) maintenance methylation and c) *de novo* methylation and assign a probability for the various possible transitions between states. In addition, we assume that there is no active demethylation (selection of appropriate targets in *A. thaliana* where this criterion holds is discussed in a later section below). In *met1* mutants, maintenance is completely compromised, whereas as in the various *h1ddm1* mutants, it is only partially compromised. We therefore consider these two cases separately beginning with the *h1ddm1* mutants.

Replication occurs in a well-defined and relatively narrow time-window of the cell cycle. For the purposes of our model, we define replication to be the start of a new cell cycle, as this is the point that new DNA is synthesized. Instead of modelling the top and bottom strands of DNA specifically, we make the standard assumption that 50% of hemi-methylated sites become unmethylated upon replication:

a) Replication:

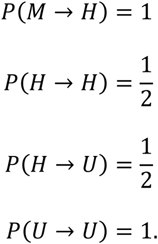

After replication, maintenance methylation by MET1 occurs semi-conservatively such that *H* → *M*. We assume that maintenance occurs both rapidly and very efficiently after replication. Indeed, for all various *h1ddm1* mutants, we find efficient maintenance to hold self-consistently (see below). We therefore re-write *r_maint_*, the contribution from the MET1-maintenance pathway, as *r_maint_* = 1 − *∊* where 0 ≤ *∊* ≪ 1 represents the probability of maintenance failure. This gives:

b) Maintenance:

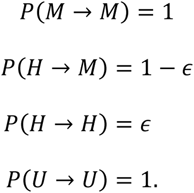

Finally, we consider the *de novo* pathways and define *P*(*U* → *H*) = 2 *r_novo_* = δ, where the factor of 2 is included as there are two possible cytosine targets in each CG site. In the mutants that we study, 0 < δ ≪ 1 is also found to hold self-consistently (see below). We neglect the possibility that *de novo* methylation could occur twice at the same CG site (i.e., a *U* → *M* transition) within a single cell cycle, as this process is of order δ^2^. We also neglect the possibility that a *de novo* event could follow a maintenance failure, as this composite process is of order εδ. The above assumptions are consistent with *de novo* methylation happening either concurrently with maintenance, or as an extended process throughout the cell cycle.

c) *De novo*:

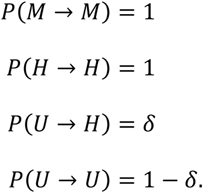

Combining the above three processes then provides the recursion relations:

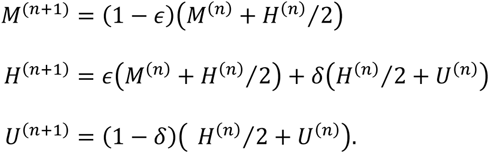

Using *U*^(*n*)^ + *H^(n)^* + *M*^(*n*)^ = 1 to eliminate *H^(n)^* gives:

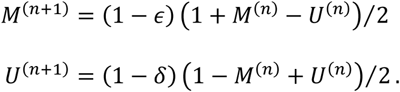

This pair of equations can be rescaled and added to provide:

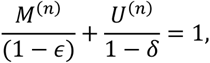

which allows the recursion relations to be expressed in terms of a single variable only, where we retain only terms linear in εand δ:

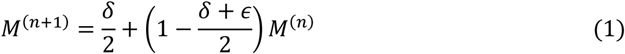

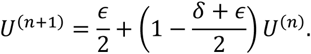

Next, we consider the *met1* mutation. The above derivation is unchanged for replication. We assume complete maintenance failure, so that *∊* = 1. As previously, we neglect processes of order δ^2^ within a single cell cycle, and later find that 0 < δ ≪ 1 also holds self-consistently for *met1* mutants. Applying these modifications provides the following recursion relations for *n* > 0:

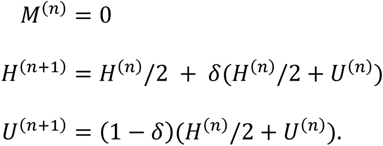

Using *H^(n)^* + *U^(n)^* = 1 then gives:

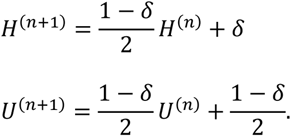

### Mathematical model for TE methylation dynamics: steady state solutions

At steady state, we define *M*^(*n*+1)^ = *M^(n)^* = *M*^∗^, and equivalently for *H* and *U*. For the various *h1ddm1* mutants, we then find that to lowest order

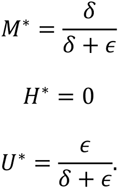

For all mutants considered here, except for *met1*, the values we extract from the data indeed give *∊,δ* ≪ 1, thereby self-consistently justifying the assumptions made in our derivation. Hence,

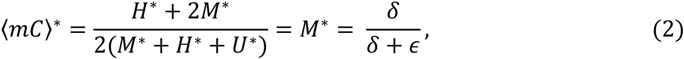

Whereas, for *met1* mutants, to lowest order:

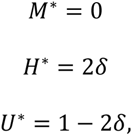

for which,

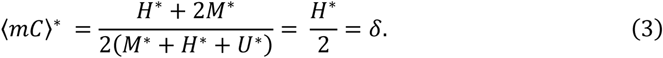

### Mathematical model for TE methylation dynamics: time dependent solutions

Using a standard approach to solve difference equations, the recursion relation in Eq. (1) can be solved to express *M^(n)^* as a function δ, *∊* and the initial fraction of methylation, *M*^0^:

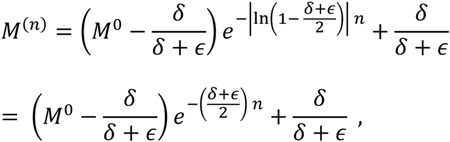

for 0 < δ ≪ 1 and 0 < *∊* ≪ 1, and therefore where we again retain only leading order terms. To allow comparison with the experimentally measured methylation level, the above exponential decay must be expressed as a function of the number of plant generations, *x*, and the number of cell cycles through the germline per plant generation, such that *n* = *n_cc_x*. The experimentally measured methylation level then becomes:

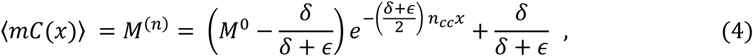

with the free parameters: δ, the *de novo* methylation rate for each unmethylated CG site per cell cycle, *∊*, the maintenance failure rate for each CG site per cell cycle, and *M*^0^, the initial methylation level before the system was perturbed away from steady state.

### Mathematical model for TE methylation dynamics: gain and loss rates per cell cycle

The overall methylation gain-rate per cell cycle (to first order) is:

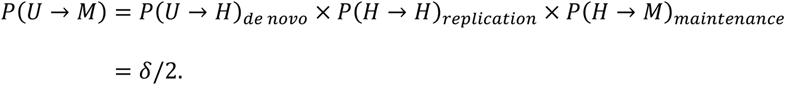

This is referred to in the main text as the ‘*de novo* rate’.

Similarly, the overall loss-rate per cell cycle (referred to in the main text as the ‘maintenance failure rate’ to emphasize that this is a passive loss of methylation) is:

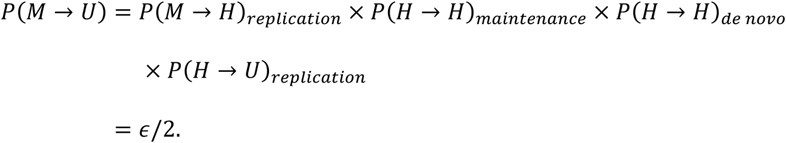

### Relationship between the mathematical model and bisulfite sequencing data from leaf tissue

Our model is applied to individual heterochromatic TEs with a length > 2 kb. As the model assumes the absence of active demethylation, we wish to exclude possible ROS1 targets from our analysis. These are known to have depleted methylation levels in *WT* plants (Lister et al., 2008; Tang et al., 2016; Zhu et al., 2007). Therefore, we select only those TEs with a *WT* methylation level of *M_WT_* > 0.65 for both replicates. This results in 1883 TEs which we fit using the model.

The transgenerational bisulfite sequencing data of leaf tissues show decreasing methylation levels as a function of time for the following mutants: *h1ddm1*, *h1ddm1cmt2*, *h1ddm1cmt3* and *h1ddm1drm2*. We fit these methylation timeseries to the decaying exponential in Eq. (4) (the numerical details are explained in the following section). Consequently, we need to know the effective timepoints at which the leaf-tissue sequencing data is obtained, as described and illustrated schematically below.

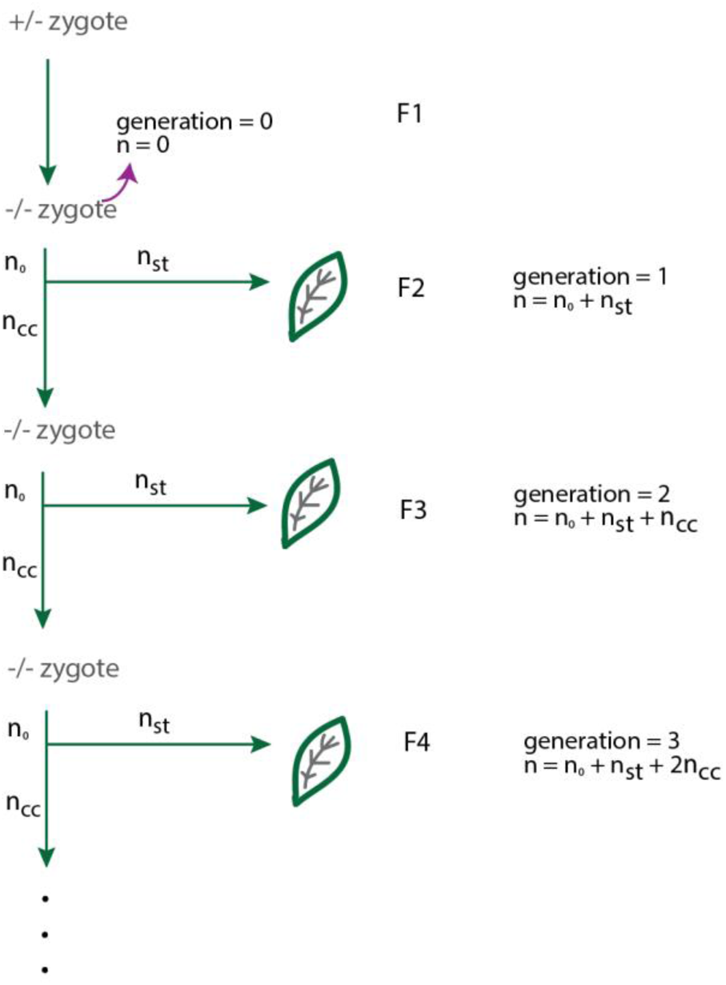

We assume that the heterozygous plant methylation level is at its steady state and define *n* = 0 to occur when the homozygous offspring is formed, coinciding with (we assume) an abrupt potential change in the values of δ and *∊*. The changed δ and *∊* values will now correspond to a different steady-state methylation level. The model predicts that the methylation level of the mutant plant will decay exponentially with time towards this new value. To fit the experimental data to this exponential decay, we first need to know the approximate number of replication events between data points.

Following (Watson et al., 2016), we assume a constant number of cell divisions through the cell lineage leading to the generative lineage (“germline”), *n_cc_* = 34. We also assume a constant number of somatic cell divisions to produce the sequenced leaf tissue: *n_st_*, this value, however, is unknown. Under these assumptions, for plants of generation F2 and later, there is a uniform number of cell divisions between each generation of Δ*n* = *n_cc_*. The heterozygous plant, however, is more problematic as it is assumed to stay at its original steady-state methylation level up until the formation of the homozygous zygote. The F2 generation is therefore separated by Δ*n* = *n*_0_ + *n_st_*, where n_0_ is the number of replications before the leaf tissue branches from the germline, both values of which are unknown.

The data points for early generations are particularly valuable as the methylation level changes most rapidly between these. We treat biological replicates of the same generation as multiple data points with the same time-value. As an initial test of the model, we first fit to the methylation level averaged over all the selected TEs (Fig. 5B). Here all generations from F2 onwards are included. We find excellent fits to both the *h1ddm1* and the *h1ddm1drm2* datasets, along with a good fit for *h1ddm1cmt2*. However, in the latter case, the replicates have a greater spread, increasing the fit uncertainty. Furthermore, for *h1ddm1cmt3*, the important F2 generation is not available significantly reducing our confidence in this fit. To partly overcome this difficulty, we use our other data sets to generate an extra effective datapoint corresponding to the *WT* plant. As demonstrated by the fit to the average mCG level for *h1ddm1* F2 to F11 replicates (fit without *WT*: Fig. 5B solid line), the mCG level clearly decays from a high initial value at *x* = 0 (*σ_fit_* = 0.011). By projecting the *WT* leaf methylation level onto the *h1ddm1* fit without WT, we find (coincidentally) that the *WT* leaf methylation level corresponds to *x* = 0 (to within the fit uncertainty). For each individual TE, therefore, we approximate the initial methylation level (at *x* = 0) by the two *WT* leaf replicates and confirm that the fit to average methylation levels when including these two extra effective datapoints (fit with *WT*: Fig. 5B dashed line) is highly consistent to the fit without *WT*. We then fit to the methylation timeseries for individual TEs to generate distributions of *δ* and *∊* values for each mutant (described in detail below). We confirmed that fits excluding the *WT* leaf replicates at *x* = 0 provide comparable distributions, however, manual inspection revealed these to provide less reliable estimates of the decay constant than the fits with the *WT* leaf replicates included (due to some fits generating anomalously low initial methylation levels and consequently unreliable values for the decay constant). All presented fits, therefore, include the *WT* replicates at *x* = 0 unless otherwise stated. For *met1* mutants the methylation decay is too rapid to be accessible in the transgenerational data. Only the steady-state methylation level is available, which corresponds directly to δ (see Eq. (3)).

### Mathematical model for TE methylation dynamics: numerical fits

The methylation timeseries, 〈*mC*〉(*x*), are fit using *curve_fit* from the *optimize* library in *scipy* (Virtanen et al., 2020). First, we performed fits using three free parameters: *M*_0_, *M*^∗^ (both as defined previously), and *δ*, the decay constant:

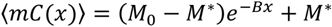

and the total fit uncertainty, **σ*_fit_*, is estimated using the sum of the diagonal of the covariance matrix:

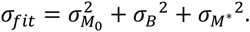

The fit parameters were constrained to 0 < **M**_0_ < 1, 0 < **M**^∗^ < 1 and *B* > 0, while using initial values of: *M*_0_ = 0.8, *B* = 1 and *M*^∗^ = 0.3. We do not fit to δ and *∊* directly, as the numerical routine is more reliable when fitting to parameters with an order of magnitude close to one. As *B* = n*_cc_* + δ*∊*)/2, we reject any fits for which *B* > *n_cc_* as this corresponds to an unphysical value for δ or *∊*. The majority of TEs produce good fits and a small minority give exceptionally poor fits. To be conservative, we reject all fits with *σ_fit_* > 1. A reliable estimate of the decay constant requires an appreciable decrease in the methylation level with time. We therefore exclude TEs with ⟨*mCG*⟩_{*F*2,*F*3}_ ≤ *λM*∗ for *h1ddm1*, *h1ddm1cmt2* and *h1ddm1cmt3* or ⟨*mCG*⟩_{*F*3,*F*4}_ ≤ *λM*^∗^ for *h1ddm1cmt3* using a threshold of *λ* = 1.05, where ⟨*mCG*⟩_{*F*2,*F*3}_ is the average methylation level of all F2 and F3 replicates for a particular mutant. Upon manual inspection of the remaining fits, we also excluded nine further TEs. Finally, we selected the subset of TEs with acceptable fits for all four mutants (amounting to 1617 TEs, over 85% of the initial set that we attempted to fit).

The values of δ and *∊* are then given by:

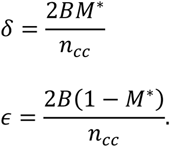

For a function *f*(*x*_1_, *x*_2_, *x*_3_) the uncertainty in *f* is given by

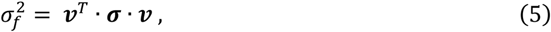

where σ is the covariance matrix for *x*_1_, *x*_2_ and *x*_3_ and ν is a vector with elements 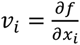. The uncertainties in the fitted values of δ and *∊* are therefore:

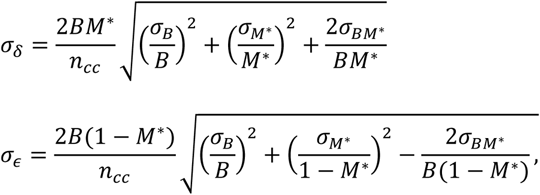

where *σ_BM∗_* is the covariance of *B* and *M*^∗^.

Two-sided paired t-tests were performed for all mutant pairings of *∊*-distributions (p-value < 1 × 10^−170^ for all pairings) and, similarly, for all mutant pairings of δ-distributions p-value < 1 × 10^−200^ for all pairings). Finally, we note that the presence of the small peak at low δ-values (Fig. 7F) is likely an artifact arising for TEs with a small decay constant. Due to their slow approach to steady-state, the fit algorithm may underestimate the precise value of *M*^∗^, and hence δ.

### Mathematical model for TE methylation dynamics: limitations of the model

We have assumed that the *de novo* rate, *δ*, is independent of the existing methylation level. This may not always be the case. For example, a decrease in RdDM targeting as more methylation is lost is possible, as appears to be the case for a small subset of TEs (Fig. S6F-H). Alternatively, there may be an analogous cooperative *de novo* pathway to that proposed for clustered CG sites as found in CG islands of mammalian genomes (Lövkvist et al., 2016), in which case δ would represent the average *de novo* rate across the whole TE. Due to the cooperativity, the effective δ could then reduce across generations as the overall methylation level declines. In the case of small variations of the *de novo* rate over successive generations, the model will still produce a good fit to a decaying exponential with effective rates for both δ and *∊*. A much larger variation in δ as a function of time, however, would likely result in a clearly visibly non-exponential decay of the methylation level, which is not observed in our data.

A further issue is that the values of δ and *∊* could each be different in the germline versus in somatic tissue (e.g., in the leaf). Our model assumes that these values are the same. Experimental data indicates that F5 mCG levels in the *h1ddm1* leaf and sperm are similar (Fig. 2A), which supports similar values of δ and *∊* in the germline and somatic tissue.

The model also assumes that all CG sites within a given TE are equivalent and all exhibit the same dynamical turnover of methylation, parameterized through δ and *∊*. However, as is apparent from the per-CG site methylation histograms (Fig. 1F, S1H), a minority of CG sites are always highly methylated or always exhibit a near-absence of methylation, configurations that are essentially excluded within our model. Future model developments may need to better incorporate this heterogeneity.

### Use of previously published datasets

GSE179796: ChIP-seq data from (Choi et al., 2021).

GSE122394: *met1* and *h1met1* bs-seq data from (Choi et al., 2020).

GSE96994: *h1* bs-seq and nucleosome data from (Lyons & Zilberman, 2017).

GSE150436: H2A.W ChIP, *ddm1* x Col-0 bs-seq, and RNA-seq from (Osakabe et al., 2021).

DRS019614: *ddm1* G9 (F10) bs-seq data from (Ito et al., 2015)

(https://ddbj.nig.ac.jp/resource/sra-sample/DRS019614).

## Author contributions

D.B.L. conceived the research plan, performed experiments and analysis, and wrote the paper

A.B. developed the mathematical model, performed analysis, and wrote the paper

S.H. performed sperm isolation and library preparation

J.Choi contributed to RNA-seq analysis

E.H. contributed to DNA methylation analysis

J. Colicchio measured plant phenotypes

I. A. measured plant phenotypes

X.F. supervised experiments on sperm DNA methylation

M.H. developed the mathematical model, supervised analysis, and wrote the paper

D.Z. conceived the research plan, supervised experiments, and wrote the paper

## Acknowledgements

The authors would like to thank Lesley Philips, Timothy Wells, and Sophie Able from Horticulture Services at the John Innes Centre. This work was supported by a European Research Council grant MaintainMeth (725746) to D.Z. D.B.L. was supported by a postdoctoral fellowship from the Helen Hay Whitney Foundation.

## Supplemental Figure Legends

**Supplemental Figure 1.**
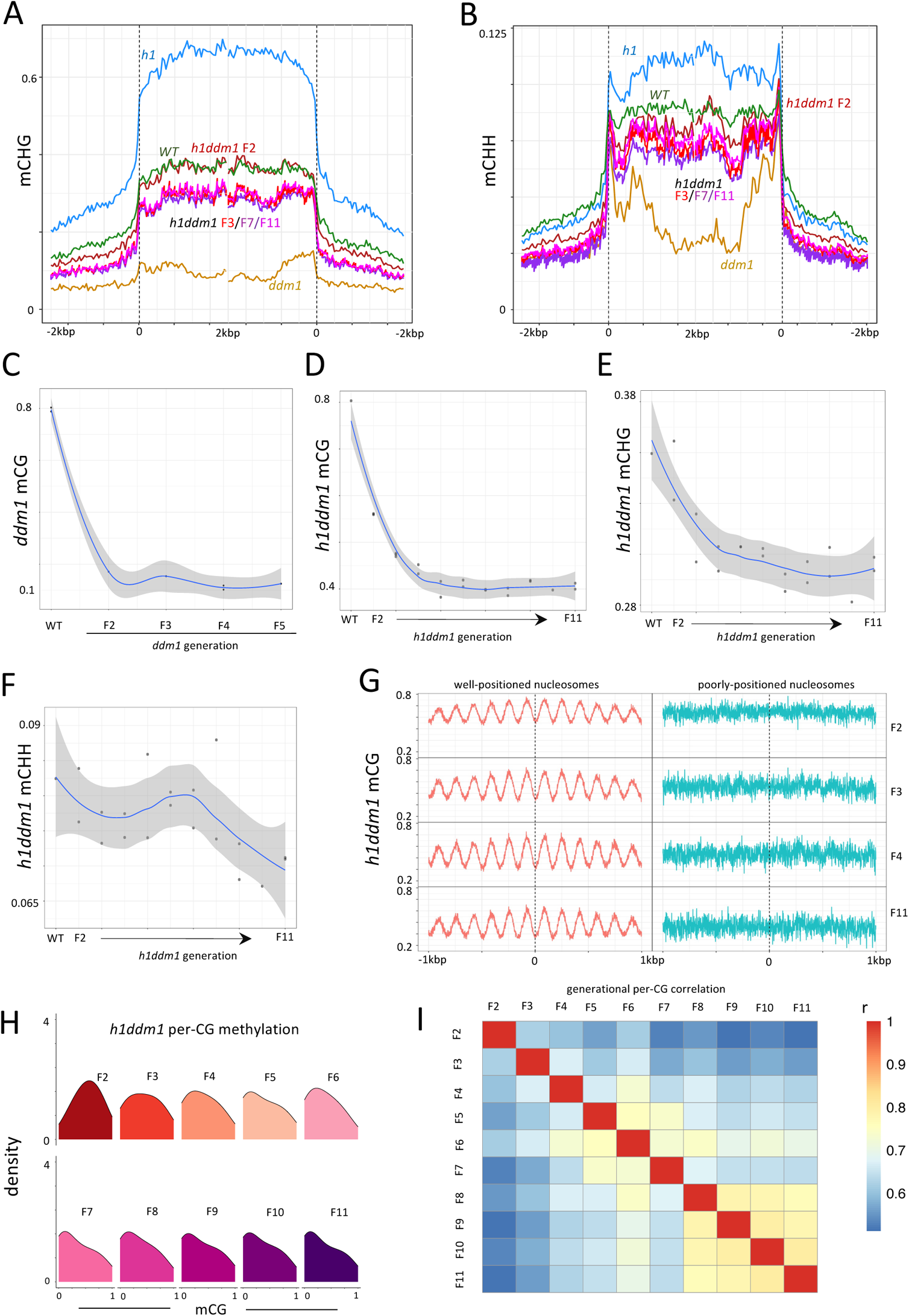
**A-B**) Average fractional *h1ddm1* mCHG (**A**) and mCHH (**B**) is plotted for all heterochromatic TEs >250bp for indicated genotypes around TE start and stop sites, as in Fig. 1A. **C**) Average *ddm1* mCG per generation (per replicate) from F2, beginning with *WT*, to highlight the steep drop in mCG that does not match that of the gradual *h1ddm1* pattern (note differences in y-axes values with panel **D**). Points are overlaid with a LOESS curve, smoothed with default ggplot stat_smooth settings (span=0.75, polynomial degree=2), with 95% confidence interval shown in grey. **D-F**) Average *h1ddm1* mCG (**D**), mCHG (**E**), and mCHH (**F**) in heterochromatic TEs plotted as single points per generation (per replicate) overlaid with a LOESS curve as in (**C**). **G)** Average *h1ddm1* mCG at well-positioned (red, left panels) or poorly-positioned (cyan, right panels) nucleosomes in heterochromatic TEs (Lyons & Zilberman, 2017). Generation of *h1ddm1* methylation data is indicated on the right. **H)** Density plots of per-CG methylation for all CG sites with coverage > 10 in heterochromatic TEs in all *h1ddm1* generations assayed. **I)** Pairwise per-TE mCG Pearson’s correlations for indicated *h1ddm1* generations represented as a heatmap.

**Supplemental Figure 2.**
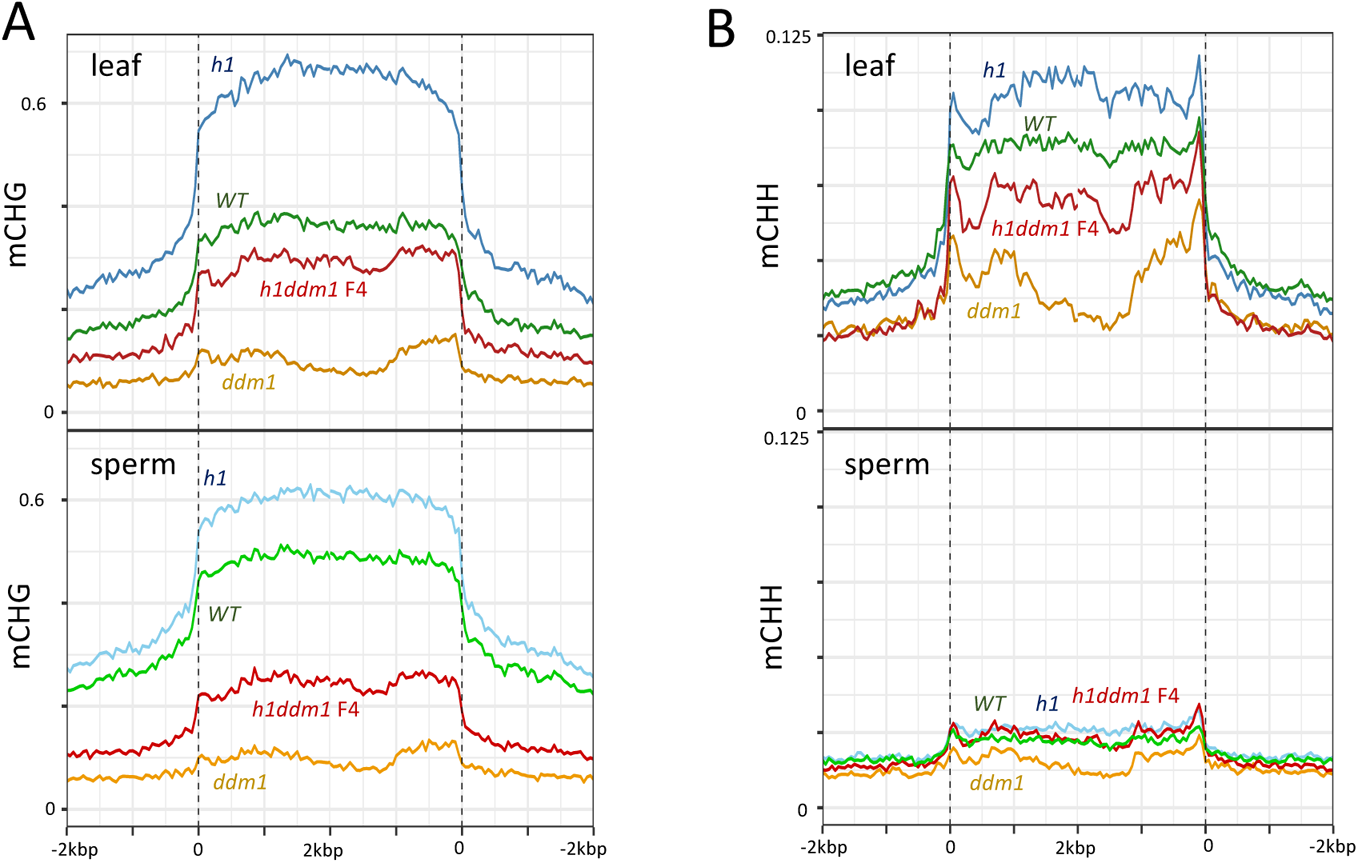
**A-B**) Line plots of average *h1ddm1* mCHG (**A**) and mCHH (**B**) at all heterochromatic TEs for the indicated genotypes around TE start and stop sites. Leaf is shown at the top, sperm at the bottom.

**Supplemental Figure 3.**
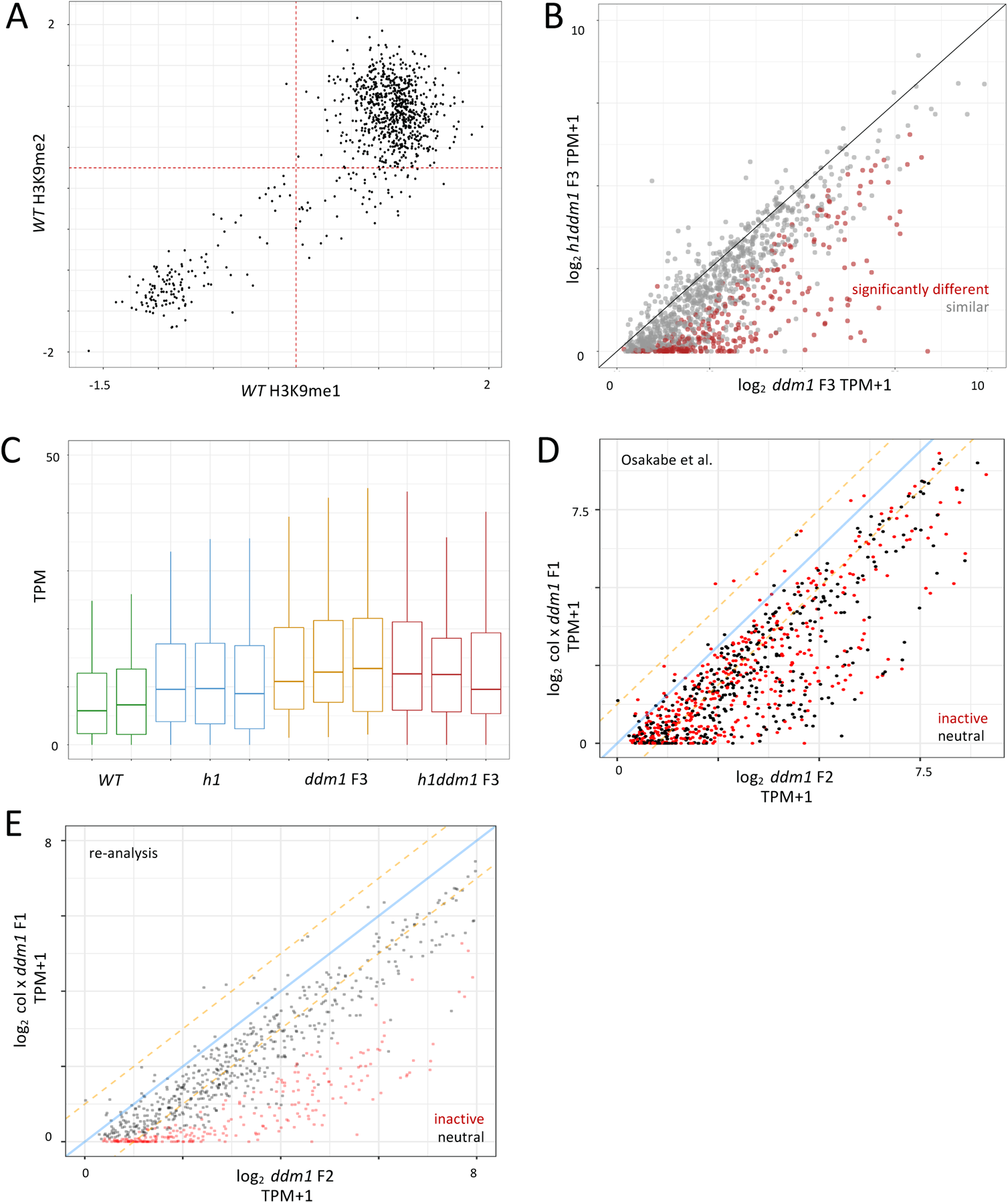
**A)** Scatterplot of *WT* H3K9me2 vs. *WT* H3K9me1 at *ddm1* transcribed loci illustrating that most upregulated *ddm1* transcripts are in regions with heterochromatic histone modifications. **B)** Scatterplot of *ddm1* vs. *h1ddm1* average expression levels for heterochromatic transcripts. Color indicates whether transcript was found to be significantly changed (q-val <0.05, likelihood ratio test per (Pimentel et al., 2017)) in *h1ddm1*. **C)** Boxplots of expression levels of euchromatic loci upregulated in *ddm1* for the indicated genotypes, with biological replicates shown individually. **D)** Scatterplot of Col-0 x *ddm1 vs. ddm1* F1 average expression levels for TE genes as defined in (Osakabe et al., 2021) using TPM from reanalysis performed in this study. Color indicates classification of TE gene as inactive (red) or neutral (black) based on source data from (Osakabe et al., 2021). **E)** Scatterplot of Col-0 x *ddm1* vs. *ddm1* F1 average expression levels for TE genes using TPM from reanalysis performed in this study. Red color (inactive) indicates that a transcript is both significantly downregulated at qval <0.05 and (*ddm1* / Col-0 x *ddm1* F1) >4.

**Supplemental Figure 4.**
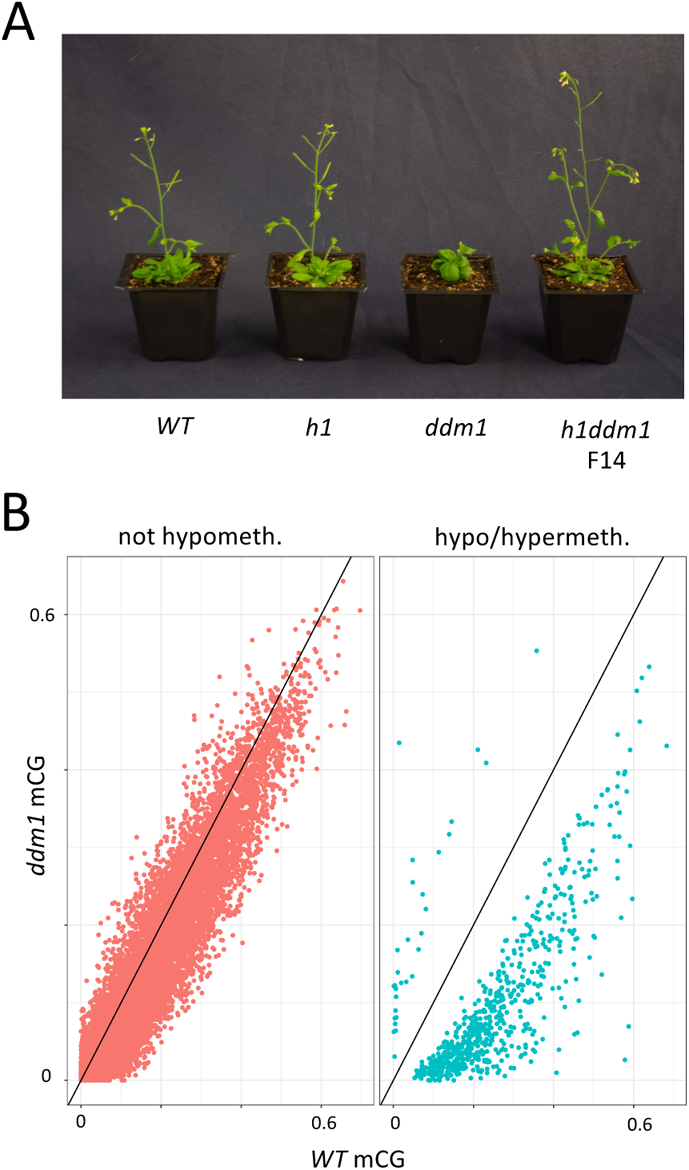
**A)** Plants at 4 weeks post-germination, genotypes are indicated. All genotypes shown are F4, except *h1ddm1* is F14. **B)** Scatterplot of genic mCG in *ddm1* vs. *WT* illustrating genes that are significantly hypomethylated in *ddm1* (n=529, right panel; p-val<1 x 10^−13^, Fisher exact test).

**Supplemental Figure 5.**
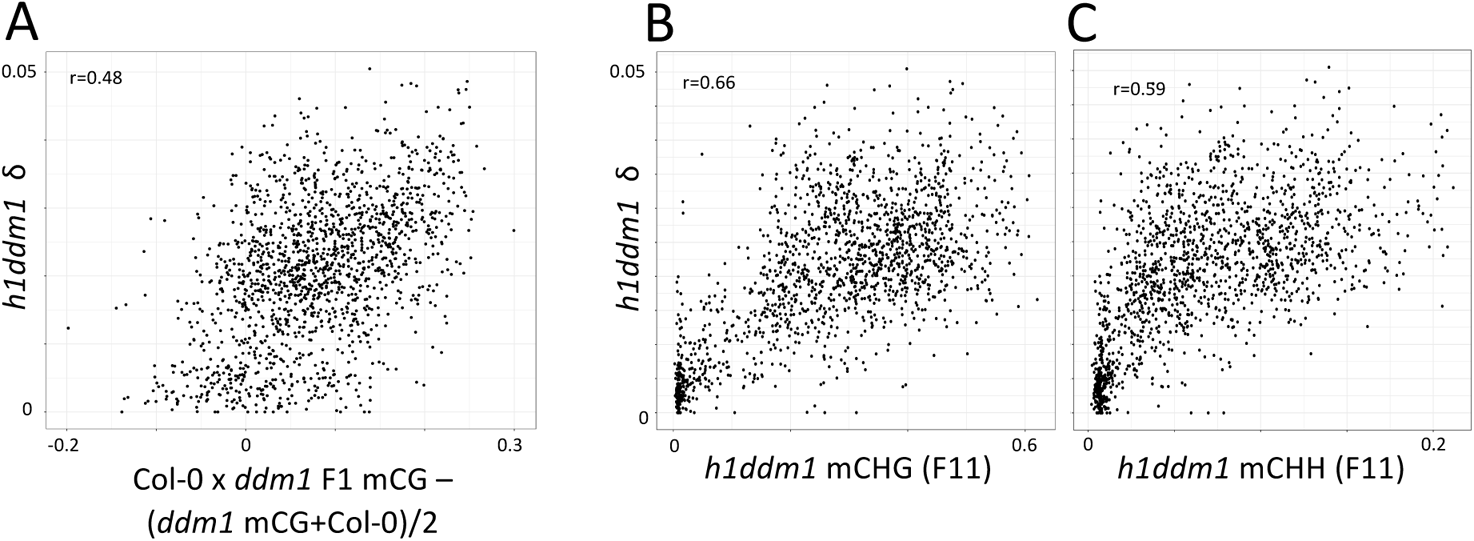
**A**) Scatterplot of *h1ddm1* δ vs. mCG in *ddm1* x Col-0 F1 as compared to average mCG of *ddm1* and *WT* (the expected level if mCG of the *ddm1* chromosomes is unchanged from *ddm1* levels in *ddm1* x Col-0 F1) at TE genes. mCG is calculated as indicated on the x-axis. **B-C**) Scatterplots of *h1ddm1* δ as a function of F11 mCHG (**B**) and mCHH (**C**). Pearson’s r is shown for both plots.

**Supplemental Figure 6.**
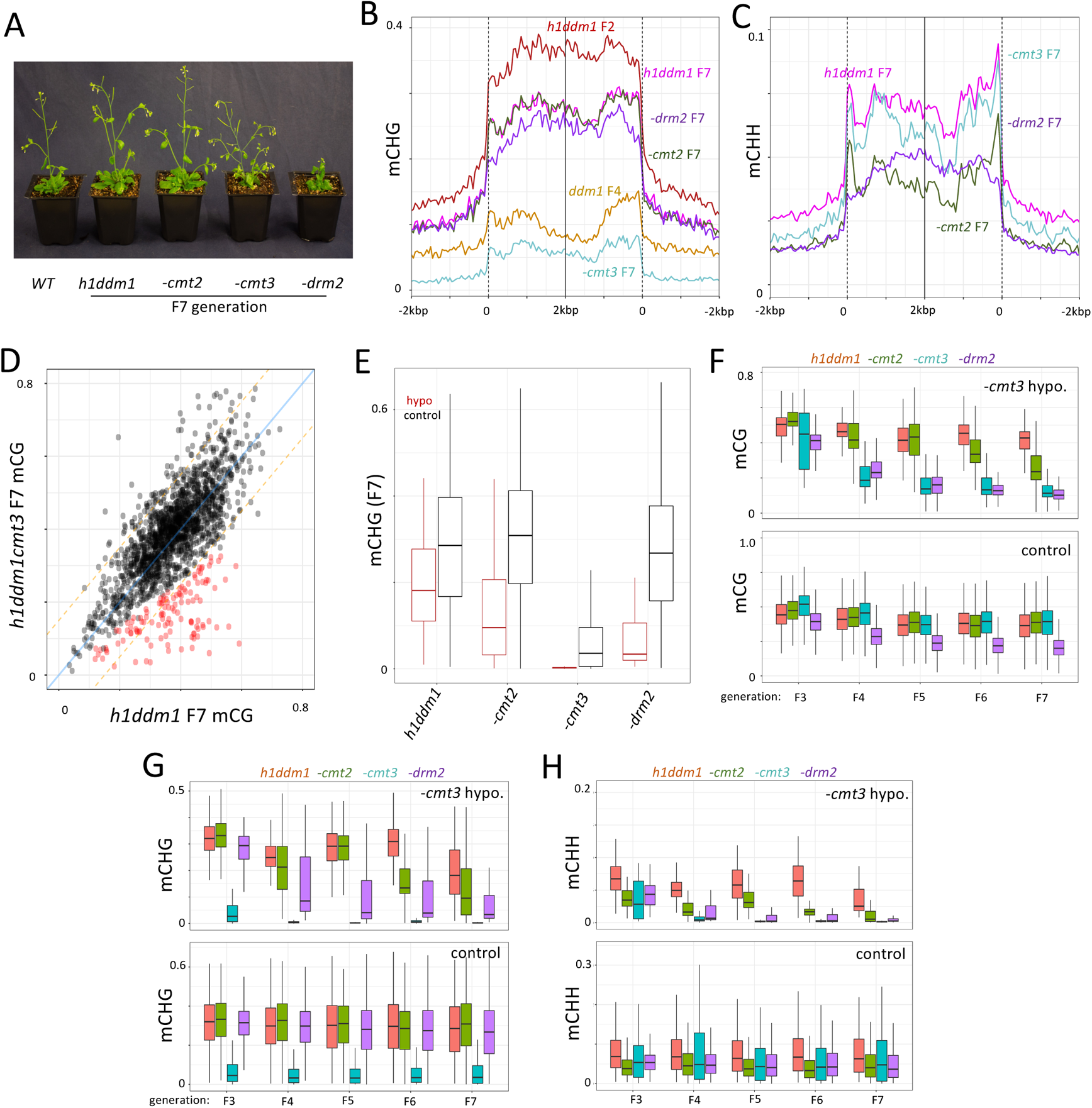
**A**) Plants at 4 weeks post-germination, genotypes are indicated such that the rightmost 3 samples are also *h1ddm1* in addition to the indicated genotype. Generation of mutants: F7; WT is F4. **B-C**) Average mCHG (**B**) and mCHH (**C**) is plotted for all heterochromatic TEs for the indicated genotypes and generations around TE start and stop sites. These were selected to illustrate depletion of mCHG in the case of *h1ddm1cmt3* and the depletion of mCHH in *h1ddm1drm2* and *h1ddm1cmt2*. **D)** Scatterplot depicting per-TE mCG of modeled TEs in *h1ddm1cmt3* vs. *h1ddm1* at F7. Red color indicates TEs significantly hypomethylated in *h1ddm1cmt3* (n=86, >50% reduction of mCG and p-val<0.01, Fisher exact test). **E)** Boxplots comparing mCHG in modeled TEs that are either hypomethylated in *h1ddm1cmt3* (red) or control TEs that are not significantly hypomethylated (black). **F-H**) Boxplots of mCG (**F**), mCHG (**G**) and mCHH (**H**) across the generations in TEs either mCG hypomethylated in *h1ddm1cmt3* (top) or all remaining TEs (bottom).

**Supplemental Figure 7.**
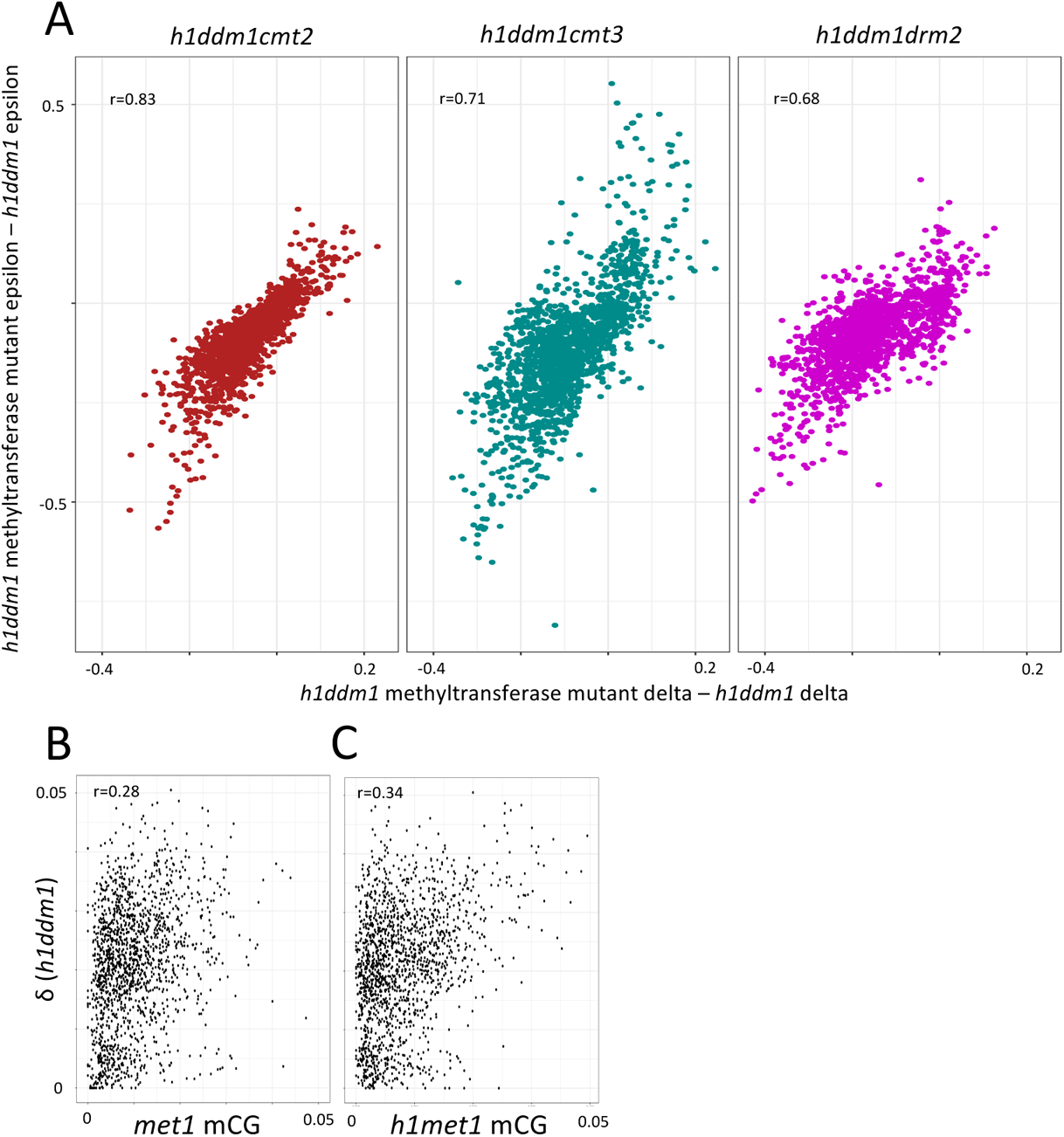
**A**) Scatterplots of changes in δ vs. changes in ε for the indicated genotype relative to *h1ddm1* values. Pearson’s r is shown. **B-C**) Scatterplots of *h1ddm1* δ vs. mCG (implied δ) in *met1* (**B**) and *h1met1* (**C**) at modeled TEs. Pearson’s r is shown.

